# FOSL2 regulates gonadotropin-dependent folliculogenesis through feedback amplification of FSH/FSHR signaling

**DOI:** 10.1101/2025.03.11.642212

**Authors:** Hongru Shi, Chaoli Chen, Zaohong Ran, Jianning Liao, Zian Wu, Xiaodong Wang, Yongheng Zhao, Wenkai Ke, Bowen Tan, Yun Liu, Youqiang Su, Wei Ren, Xiang Li, Changjiu He

## Abstract

The meticulous orchestration of gonadotropin-dependent folliculogenesis constitutes the cornerstone of female reproductive cyclicity and fertility, with FSH/FSHR signaling recognized as the master regulator. Achieving the necessary amplification of this signaling is essential for successful GTH-dependent folliculogenesis, yet the mechanisms remain inadequately defined. Our study utilizes single-cell and spatial transcriptomics to identify FOSL2 as an FSH-inducible transcription factor, exhibiting precise spatiotemporal co-expression with FSHR. FOSL2 knockdown in vitro resulted in notable reductions in granulosa cell proliferation, induced apoptosis, and disrupted gonadotropin-dependent folliculogenesis. In vivo studies using conditional FOSL2 deletion in mouse granulosa cells corroborated these results, demonstrating a complete halt in GTH-dependent folliculogenesis and resultant infertility. Mechanistic exploration unveiled that FSH/FSHR initiates FOSL2 expression via the cAMP/PKA/CREB cascade, while FOSL2 in turn enhances FSHR transcription through direct promoter binding, thereby establishing a self-amplifying loop. This loop represents a molecular switch, modeled to the all-or-nothing dynamics of GTH-dependent folliculogenesis. The evolutionary conservation of this mechanism was confirmed through cross-species analyses in sheep, where FOSL2 deficiency similarly attenuated FSH/FSHR signaling and inhibited follicular growth. Our findings advance the understanding of folliculogenesis by revealing a novel FOSL2-centered amplification loop for FSH/FSHR signaling, highlighting the indispensable role of FOSL2 in reproductive biology.

## INTRODUCTION

Subfertility and infertility represent significant global health concerns, with defective folliculogenesis identified as a critical etiology [1]. Folliculogenesis commences with primordial follicle activation, and progresses through primary, secondary, small antral and big antral stages prior to ovulatory competence acquisition. A critical developmental milestone occurs with the emergence of the antrum within the follicle: pre-antral folliculogenesis is driven by gonadotropin hormone (GTH)-independent mechanisms, whereas subsequent GTH-dependent folliculogenesis requires precise regulation by follicle-stimulating hormone (FSH) and luteinizing hormone (LH) [2–4]. This GTH-dependent folliculogenesis is essential for various reproductive functions, including the initiation of the estrous cycle, estrogen-mediated maintenance of secondary sexual characteristics, sexual receptivity, and luteal formation post-ovulation. Clinically, dysregulation of this process accounts for approximately 30% of infertility cases worldwide and is pathologically linked to prevalent reproductive disorders including menstrual irregularities, polycystic ovary syndrome (PCOS), luteinized unruptured follicle syndrome, and primary/secondary amenorrhea [5].

FSH is recognized as the master regulator in regulating regulating GTH-dependent folliculogenesis by specifically targeting mural granulosa cells (GCs) via the FSH receptor (FSHR), a G protein-coupled receptor. Upon FSH binding, FSHR predominantly activates the cAMP-PKA cascade, which orchestrates GC proliferation, anti-apoptotic protection, estrogen synthesis, and antrum expansion [6]. At the molecular level, PKA phosphorylates CREB and histone H3, initiating the transcription of CRE-containing genes [7]. For non-CRE-regulated targets, PKA utilizes alternative signaling cascade such as PI3K-AKT. Studies indicate that PKA potentiates AKT phosphorylation through an indirect mechanism [8], mediating pleiotropic regulatory effects: (i) activation of mTOR to modulate target gene transcription and translation [9,10], (ii) promotion of GC proliferation and estrogen synthesis by inhibiting GSK3β-mediated suppression of CyclinD2 and β-catenin [11], and (iii) inactivation of FOXO1/3 to relieve their inhibitory effects on proliferation and anti-apoptosis [12,13].

In parallel to PI3K-AKT, FSH/FSHR also activates the RAS-RAF-ERK cascade. PKA facilitates ERK phosphorylation by dissociating the inhibitory interaction between PTP and ERK, thereby enabling downstream gene transcription [14]. Emerging evidence also highlights the role of PKA in modulating the Wnt, NF-κB, L-type Ca²⁺ channels, and Sgk pathways [11, 15–17], all of which contribute to follicular survival and developmental progression.

FSH/FSHR signaling undergoes temporally dynamic modulation rather than stable activity throughout the GTH-dependent phase [18,19], a period lasting approximately 48 hours in mice. During the initial 12-hour period, low-intensity FSH/FSHR signaling supports preparatory cellular processes such as nucleobase synthesis, DNA replication, and mRNA processing, without inducing significant follicular morphological changes. A pivotal transition occurs between 12 and 24 hours, marked by sharp amplification in FSH/FSHR signaling. This amplification triggers rapid follicle maturation characterized by a metabolic shift to glycolysis, accelerated GC proliferation, peak estrogen synthesis, and significant antrum expansion, ultimately conferring ovulatory competence to the follicle [18]. FSH/FSHR signaling amplification is essential for GTH-dependent folliculogenesis. While the molecular drivers remain incompletely understood, transcriptional upregulation of FSHR is a key requirement. Current evidence suggests the presence of a self-amplifying loop within GCs [20], where initial low-intensity FSH/FSHR signaling induces specific transcription factors (TFs), such as estrogen receptors [21], which subsequently enhance FSHR transcription, thereby further amplifying FSH/FSHR. This loop accelerates the progression from small antral follicles to pre-ovulatory follicles. Identifying FSH-responsive TFs that drive this amplification loop therefore constitutes a pivotal strategy for elucidating the molecular basis of FSH/FSHR amplification.

Using mouse and sheep models, this study identified FOSL2, a member of the FOS family, as an FSH-inducible TF that exhibits dynamic, spatiotemporal co-expression with FSHR. Genetic ablation of FOSL2 blocked GTH-dependent folliculogenesis, resulting in the loss of estrous cyclicity and complete infertility. Mechanistic analyses revealed FOSL2 deficiency impairs FSHR transcriptional upregulation and abolishes ovarian estrogen synthesis, thereby disrupting the self-amplifying loop of FSH/FSHR signaling.

## RESULTS

### 1. FOSL2 identified as an FSH-responsive TF with spatiotemporal co-expression with FSHR

To identify the FSH-responsive TFs, we performed single-cell RNA sequencing (scRNA-seq) on ovaries harvested 48 hours post-pregnant mare serum gonadotropin (PMSG, an FSH-equivalent hormone) administration. This approach facilitated the acquisition of high-resolution transcriptional landscapes of all GCs in the ovary (Table S1). Subsequently, we conducted a customized analysis of publicly available spatial transcriptomic datasets from mouse ovaries [22], which enabled the generation of spatially informed transcriptomic atlas of GCs from large preantral (Type-5b), small antral (Type-6), and large antral follicles (Type-7/8) (Table S2). By integrating scRNA-seq data with spatial transcriptomic maps (see Materials and Methods), we reconstructed a high-resolution and spatially informed transcriptional atlas of GCs encompassing the developmental continuum from Type-5b to Type-7/8 follicles (Figure 1A, Table S3). Following evaluation, the integrated transcriptome dataset demonstrated acceptable quality with gene training scores clustering around 0.8 and AUC values reaching 0.636. Systematic analysis of this integrated dataset identified 391 TFs exhibiting upregulated expression during this follicular transition (Table S4). Applying a stringent inclusion criterion of normalized expression count > 0.5 in at least one developmental stage, we further refined the list to 19 TFs with robust expression abundances, including well-known TFs that being pivotal in folliculogenesis (Foxl2, Hmgb2, Gata6, Gata4) (Figure 1D). Among these, FOSL2 emerged as a particularly compelling candidate, as qRT-PCR analysis demonstrated that its expression in the ovary was significantly higher than in other organs (Figure 1E). Immunofluorescence staining further confirmed its predominant localization within GCs (Figure 1F).

**Figure 1.**
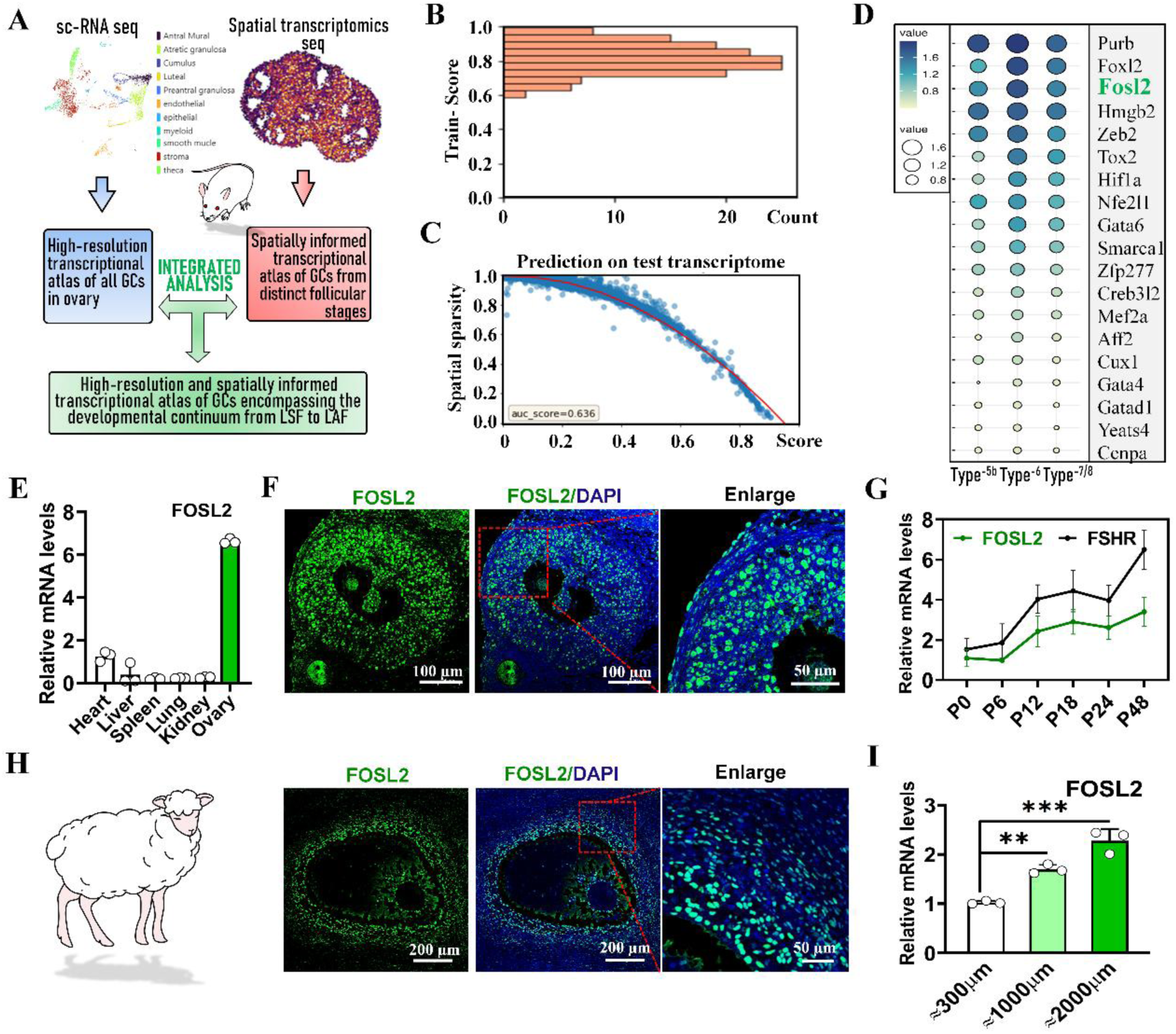
FOSL2 identified as an FSH-responsive TF with spatiotemporal co-expression with FSHR. (A) Schematic representation illustrating the integrative analysis strategy combining scRNA-seq and spatial transcriptomics. Ovaries for scRNA-seq were collected 48 hours following PMSG injected. (B) Histogram of gene similarity scores from training data. (C) Scatter plot of gene scores (x-axis: 0.0-1.0) versus spatial sparsity (y-axis: 0.0-1.0) from Tangram predictions on integrated transcriptome data. (D) TFs exhibiting high expression abundances in GCs and upregulated during transition from Type-6 to Type-7/8 as identified through integrative analysis. (E) Tissue expression profile of FOSL2 across organs. Tissues harvested 48 hours post-PMSG injection, n=3 mice. (F) Immunofluorescence localization of FOSL2 in mouse small antral follicles, with nuclear counterstaining by DAPI. (G) Time-course expression dynamics of FOSL2 and FSHR in GCs isolated at indicated timepoints post-PMSG injection, n=4 independent GC samples. (H) Immunofluorescence validation of FOSL2 localization in sheep small antral follicles, with nuclear visualization via DAPI. (I) qRT-PCR analysis of FOSL2 expression in GCs isolated from sheep follicles of different size. n=3 GC samples. Statistical significance were determined using one-way ANOVA followed by Tukey’s post hoc test, values were mean ± SD. Significant differences were denoted by **P<0.01, ***P<0.001. The experiments in panels E, G and I were repeated independently three times with consistent results.

Notably, FOSL2 expression was markedly induced following treatment with PMSG, exhibiting a dynamic spatiotemporal expression pattern that closely mirrored that of the FSHR throughout the GTH-dependent phase (Figure 1G).

Subsequent experiments analyzed FOSL2’s expression in sheep follicles, a mono-ovulatory species. Consistent with findings in mice, FOSL2 was predominantly expressed in the GCs of sheep follicles (Figure 1H), with a clear positive correlation between follicular diameter and expression levels (Figure 1I). These conserved expression dynamics across polyovulatory and monoovulatory species strongly support a regulatory role for FOSL2 in GTH-dependent folliculogenesis.

### 2. Knockdown of FOSL2 impaired GC proliferation and induces apoptosis

Next, we transfected KK1 cell line-a commercially available mouse GC line-with FOSL2-targeting small interfering RNA (siRNA) (Figure 2A), achieving efficient knockdown as validated by qPCR and Western blotting (Figure 2B, S1A). Subsequent flow cytometric cell cycle analysis revealed profound cell cycle dysregulation in FOSL2-knockdown GCs, with a marked accumulation of cells in G1 phase accompanied by concomitant reductions in S-phase and G2-phase populations (Figure 2C). This cell cycle perturbation was further corroborated by real-time cellular analysis (RTCA) showing delayed entry into exponential growth phase and reduced peak cell index values in FOSL2-knockdown GCs (Figure 2D). The proliferative defect was quantitatively confirmed through EdU incorporation assays, which demonstrated a 15.9% reduction in newly synthesized nuclei compared to control GCs (Figure 2E). FOSL2-knockdown concurrently induced apoptosis in GCs. Western blotting detected elevated levels of cleaved Caspase-3 and cleaved PARP, two hallmarks of apoptotic activation, in FOSL2-knockdown GCs (Figure S1B). Flow cytometric quantification of Annexin-V positive cells showed a significant elevation in early apoptotic populations following FOSL2 knockdown (Figure 2F). This finding was further substantiated by TUNEL staining, which revealed a significant increase in DNA fragmentation-positive nuclei compared to control GCs (Figure 2G).

**Figure 2.**
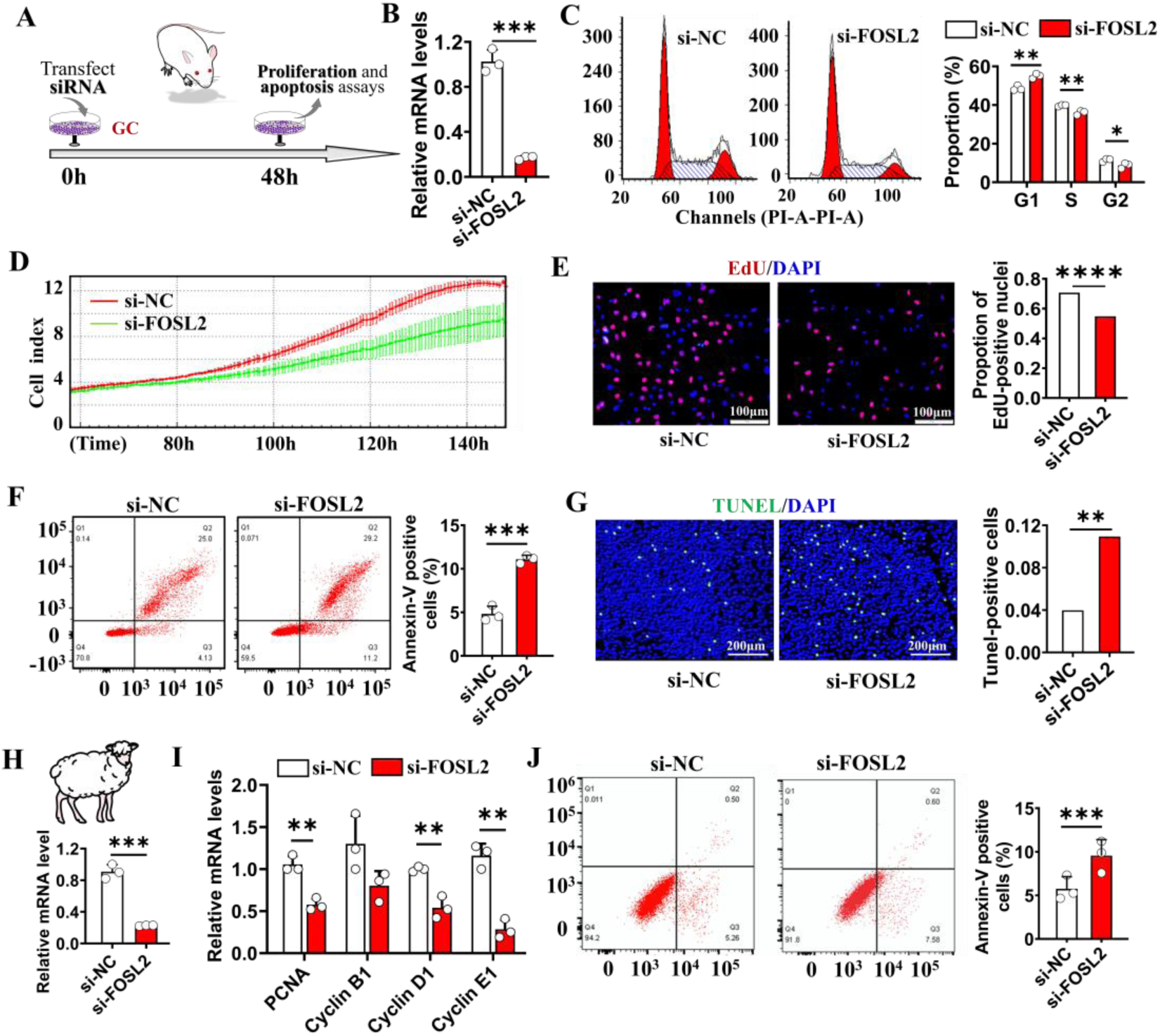
Knockdown of FOSL2 impaired GC proliferation and induces apoptosis. (A) Schematic illustration of the experimental design for FOSL2 knockdown in cultured mouse GCs. (B) Validation of FOSL2 knockdown efficiency by qRT-PCR; n = 3 independent GC samples. Scrambled siRNA (si-NC) was used as the negative control. (C) Changes in cell cycle distribution upon FOSL2 knockdown. Left: representative flow cytometry plots; right: quantification of cell cycle phase distribution. n = 3 independent GC samples. (D) Real-time monitoring of GC proliferation using the xCELLigence RTCA system. (E) Assessment of GC proliferation by EdU incorporation assay. Left: representative images showing EdU-positive cells (red). Right: quantification of EdU-positive nuclei. (F) Apoptosis analysis by flow cytometry using Annexin V staining. Left: representative dot plots; right: quantification of Annexin V-positive cells. n = 3 independent GC samples. (G) TUNEL assay for detection of apoptotic cells. Left: representative images showing TUNEL-positive cells (green). Right: quantification of apoptosis-positive cells. (H) Lentivirus-mediated shRNA knockdown of FOSL2 in sheep GCs. Knockdown efficiency was validated by qRT-PCR; n = 3 samples. GCs were isolated from small antral follicles. (I) Expression analysis of proliferation-related genes using qRT-PCR, n = 3 independent GC samples. (J) Apoptosis assessment in sheep GCs by flow cytometry with Annexin V staining. Left: representative flow cytometry plots; right: quantification of Annexin V-positive cells. n = 3 independent GC samples. Statistical significance was evaluated using a two-tailed unpaired Student’s t-test or chi-square test. Data are presented as mean ± SD. Significant differences were denoted by*P<0.05, **P<0.01 ***P<0.005, ****P<0.001. All experiments were independently repeated twice with consistent results.

To evaluate conservation of FOSL2 function, we extended analyses to sheep GCs (Figure 2H). Consistent with observations in mouse GCs, FOSL2 knockdown in sheep GCs significantly downregulated the expression of proliferation-associated markers including PCNA, Cyclin D1, and Cyclin E1 compared to control group (Figure 2I). Moreover, FOSL2 knockdown significantly increased Annexin-V positive cells as quantified by flow cytometric analysis (Figure 2J).

### 3. Knockdown of FOSL2 impaired GTH-dependent folliculogenesis in vitro

To elucidate the functional relevance of FOSL2 during folliculogenesis, we performed stage-specific lentiviral shRNA-mediated knockdown of FOSL2 in cultured ovarian follicles, aiming to dissect its role at distinct developmental stages (Figure 3A, B). In small preantral follicles, targeted knockdown of FOSL2 over a 5-day ex vivo culture period did not result in any overt morphological or functional abnormalities. Quantitative assessments revealed no significant differences in follicular size (Figure 3C), the proportion of EdU-positive GCs (Figure S2A), or the antral fraction (Figure 3D) between the knockdown and control groups. These findings suggest that FOSL2 is dispensable for preantral follicle development and antrum emergence.

**Figure 3.**
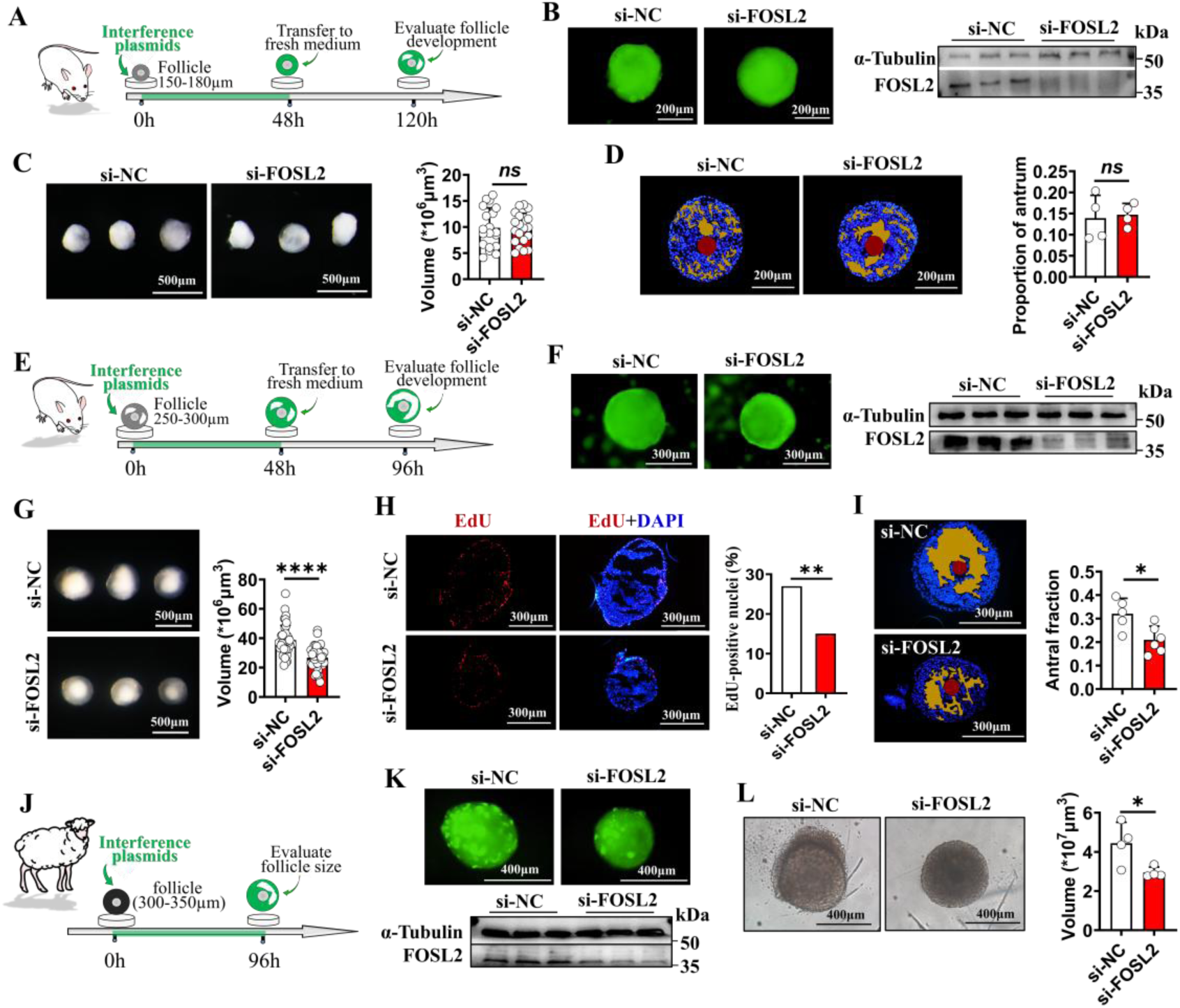
Knockdown of FOSL2 impaired GTH-dependent folliculogenesis in vitro. (A) Schematic illustration of the experimental design for FOSL2 knockdown in mouse GTH-independent follicles. (B) Evaluation of FOSL2 knockdown efficiency by Western blotting; n = 3 independent follicular samples. Green fluorescence indicates successful plasmid transfection in follicles. Original blots were provided in Figure S6. (C) Analysis of follicle volume changes following FOSL2 knockdown; n = 19 follicles. (D) Change in follicular antrum fraction following FOSL2 knockdown; n = 4 follicles. Yellow area represents the follicular antrum. (E) Schematic illustration of the experimental design for FOSL2 knockdown in mouse GTH-dependent follicles. (F) Evaluation of FOSL2 knockdown efficiency by Western blotting; n = 3 independent follicular samples. Original blots were provided in Figure S6. (G) Follicle volume changes following FOSL2 knockdown; n = 42 follicles (si-NC), n = 37 follicles (si-FOSL2). (H) Follicular cell proliferation analysis using the EdU incorporation assay. Left: representative images of EdU staining; right: quantification of EdU-positive nuclei; n = 7 (si-NC) and n = 6 (si-FOSL2) follicles. (I) Changes in the follicular antrum fraction post-FOSL2 knockdown; n = 5 (si-NC) and n = 6 (si-FOSL2) follicles. (J) Schematic illustration of the experimental design for FOSL2 knockdown in sheep GTH-dependent follicles. (K) Evaluation of FOSL2 knockdown efficiency by Western blotting; n = 3 independent follicular samples. Original blots were provided in Figure S6. Green fluorescence indicates successful transcription of interfering plasmids in sheep follicles. (L) Analysis of sheep follicle volume following FOSL2 knockdown; n = 4 follicles. Statistical significance was determined using two-tailed unpaired Student’s t test or chi-square test, values were mean ± SD. Significant differences were denoted by*P<0.05, **P<0.01 ****P<0.001. The experiments in panels C, D, G, I and L were repeated independently twice with consistent results.

In contrast, FOSL2 knockdown in small antral follicles led to notable developmental impairments. After 4 days of culture, the knockdown group exhibited a reduced follicular size (Figure 3E–G), downregulated expression of proliferation-related genes (Figure S2B), and a lower proportion of EdU-positive GCs compared to controls (Figure 3H). Additionally, knockdown follicles demonstrated defective antrum expansion, characterized by an 11.4% reduction in the antral fraction (Figure 3J). These results indicate that FOSL2 is critically involved in regulating GTH-dependent folliculogenesis.

Consistent with the phenotype observed in mouse follicles, FOSL2-knockdown sheep follicles also displayed a reduced size following 4-day culture relative to controls (Figure 3K, L), highlighting a conserved role for FOSL2 in mammalian folliculogenesis across species.

### 4. GC-specific FOSL2 knockout blocked GTH-dependent folliculogenesis and caused female infertility

To further investigate the functional necessity of FOSL2 in GTH-dependent folliculogenesis, we generated GC-specific FOSL2 conditional knockout (cKO-FOSL2) mice using an FSHR-Cre driver system (Figure 4A). Immunofluorescence staining (Figure 4B) and Western blotting (Figure 4C) assays confirmed the efficient and specific ablation of FOSL2 in ovarian GCs. Phenotypic analysis of cKO-FOSL2 females revealed significant reproductive defects. Specifically, longitudinal observation across three consecutive estrous cycles showed complete cessation of cyclic estrous behavior in cKO mice (Figure 4D). During a 15-day mating trial, cKO females failed to produce copulatory plugs and did not give birth to offspring (Figure 4E).

**Figure 4.**
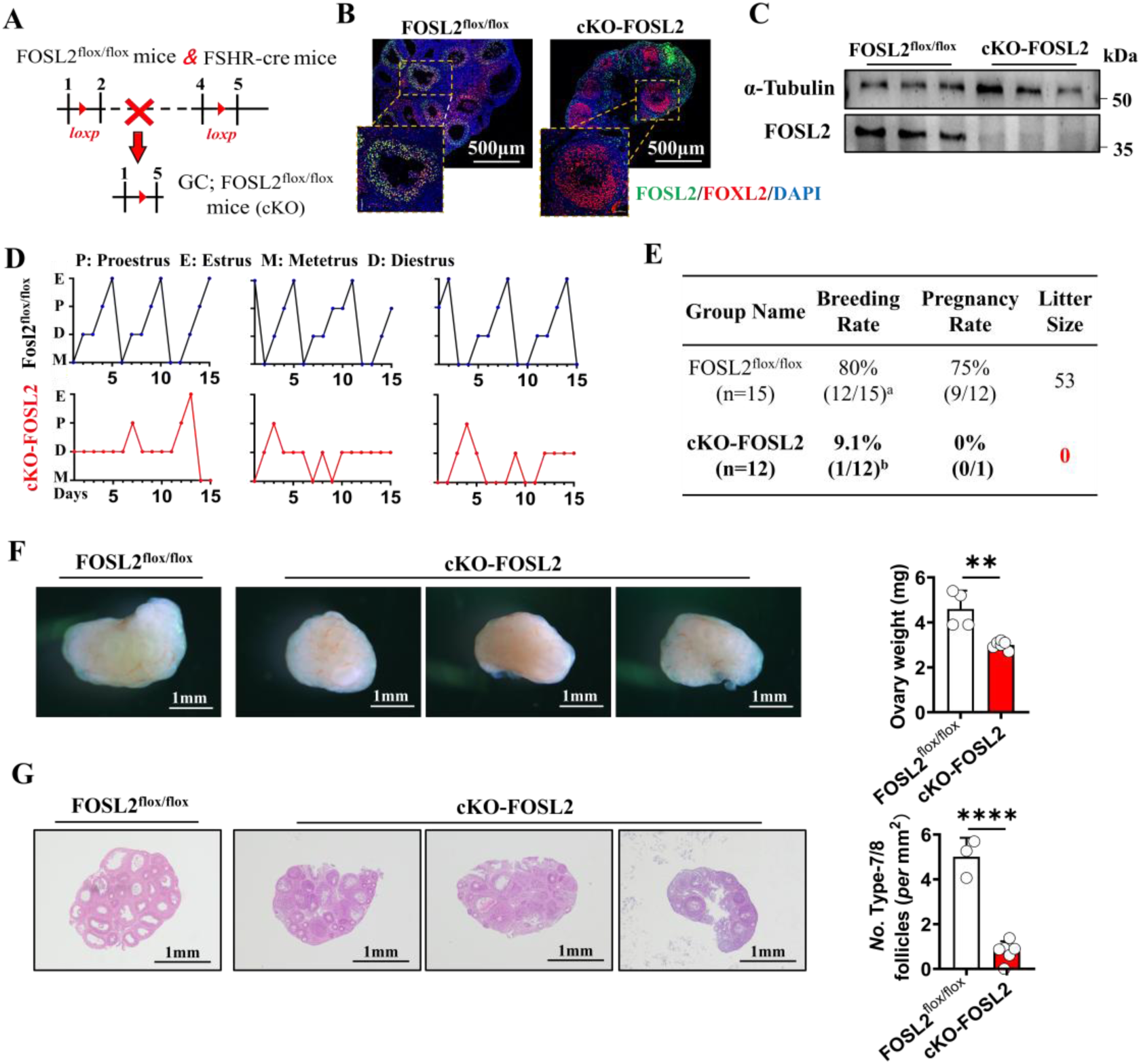
GC-specific FOSL2 knockout blocked GTH-dependent folliculogenesis and caused female infertility. (A) Schematic representation of the conditional knockout of FOSL2 in mouse GCs. (B) Immunostaining confirming successful deletion of FOSL2 in GCs. Blue: DAPI (nuclear stain), Green: FOSL2, Red: FOXL2. (C) Western blotting analysis confirming the successful deletion of FOSL2 in GCs; n = 3 independent GC samples. GCs were isolated from ovaries 48 hours post-PMSG injection. Original blots were provided in Figure S6. (D) Evaluation of estrous cycle alterations following FOSL2 knockout, n=10 mice. (E) Assessment of reproductive capacity in cKO-FOSL2 mice. (F) Analysis of ovarian weight. Left: representative images of ovaries; right: quantification of ovarian weight; n = 4 (FOSL2^flox/flox^) and 6 (cKO-FOSL2) biological independent ovaries. Ovaries were harvested 48 hours post-PMSG injection. (G) Morphological examination of ovaries using H&E staining. Left: representative images; right: quantification of Type-7/8 follicles. n=3 (FOSL2^flox/flox^) and 6 (cKO-FOSL2) biological independent ovaries. Statistical significance was determined using two-tailed unpaired Student’s t test, values were mean ± SD. Significant differences were denoted by **P<0.01 ****P<0.001. The experiments in panels F and G were repeated independently twice with consistent results.

Ovarian phenotyping performed 48 hours after PMSG administration revealed substantially reduced ovarian weights in cKO mice compared to control FOSL2^flox/flox^ littermates (Figure 4F). Histological examination revealed a developmental arrest at the small antral follicle stage in cKO ovaries, whereas control ovaries exhibited abundant Type-7/8 follicles (Figure 4G). Furthermore, GCs isolated from cKO ovaries displayed marked downregulation of proliferation-associated genes (PCNA, Ki67 and Cyclin E1) and concomitant upregulation of pro-apoptotic proteins (cleaved caspase-3, Bax) relative to FOSL2^flox/flox^ controls (Figure S3). These morphological and molecular alterations provide compelling evidence that disruption of FOSL2 function impedes GTH-dependent folliculogenesis, thereby establishing a mechanistic link between the observed reproductive phenotypes (anestrus and infertility) and impaired follicle maturation. These findings establish FOSL2 as an indispensable TF required for GTH-dependent folliculogenesis. Notably, despite the blockade of GTH-dependent folliculogenesis in cKO mice, they were still capable of ovulating approximately half the number of oocytes compared to FOSL2 ^flox/flox^ mice following the administration of 5 IU PMSG (Figure S4).

### 5. FOSL2 amplified FSH/FSHR signaling through transcriptionally reinforcing the expression of FSHR and CYP11A1

To investigate the molecular mechanisms of GTH-dependent folliculogenesis arrest in FOSL2-deficient mice, we employed bioinformatic approaches to predict FOSL2-regulated target genes. Utilizing JASPAR (http://jaspar.genereg.net/) and FIMO (https://meme-suite.org/) databases, we identified conserved FOSL2-binding cis-regulatory elements within the promoter regions of 5 genes (LHCGR, STAR, INHA, FSHR, and CYP11A1) essential for GTH-dependent folliculogenesis (Figure 5A). Validation through electrophoretic mobility shift assays (EMSA) further confirmed direct physical interactions between recombinant FOSL2 protein and promoters of FSHR, CYP11A1 and LHCGR (Figure 5B). Building on these results, we hypothesized that FOSL2 modulates GTH-dependent folliculogenesis by amplifying FSH/FSHR signaling, primarily through transcriptional reinforcement of FSHR and steroidogenic capacity. Subsequent dual-luciferase reporter assays demonstrated that FOSL2 significantly enhances the transcriptional activity of both FSHR and CYP11A1 promoters (Figure 5C). Consistently, qRT-PCR analysis revealed markedly reduced mRNA levels of FSHR and CYP11A1 in cKO-FOSL2 GCs compared to FOSL2^flox/flox^ controls (Figure 5D). Radioimmunoassay further demonstrated a significant reduction in serum estradiol levels in cKO mice compared to FOSL2^flox/flox^ control mice (Figure 5E). Finally, we quantitatively assessed the activity of three canonical FSH-downstream signaling cascades: cAMP-PKA, RAS-RAF-ERK, and mTOR. Western blotting results revealed that FOSL2-knockdown significantly suppressed the enhancement of these three cascades in GCs stimulated by FSH supplementation (Figure 5F), consistent with our expectation.

**Figure 5.**
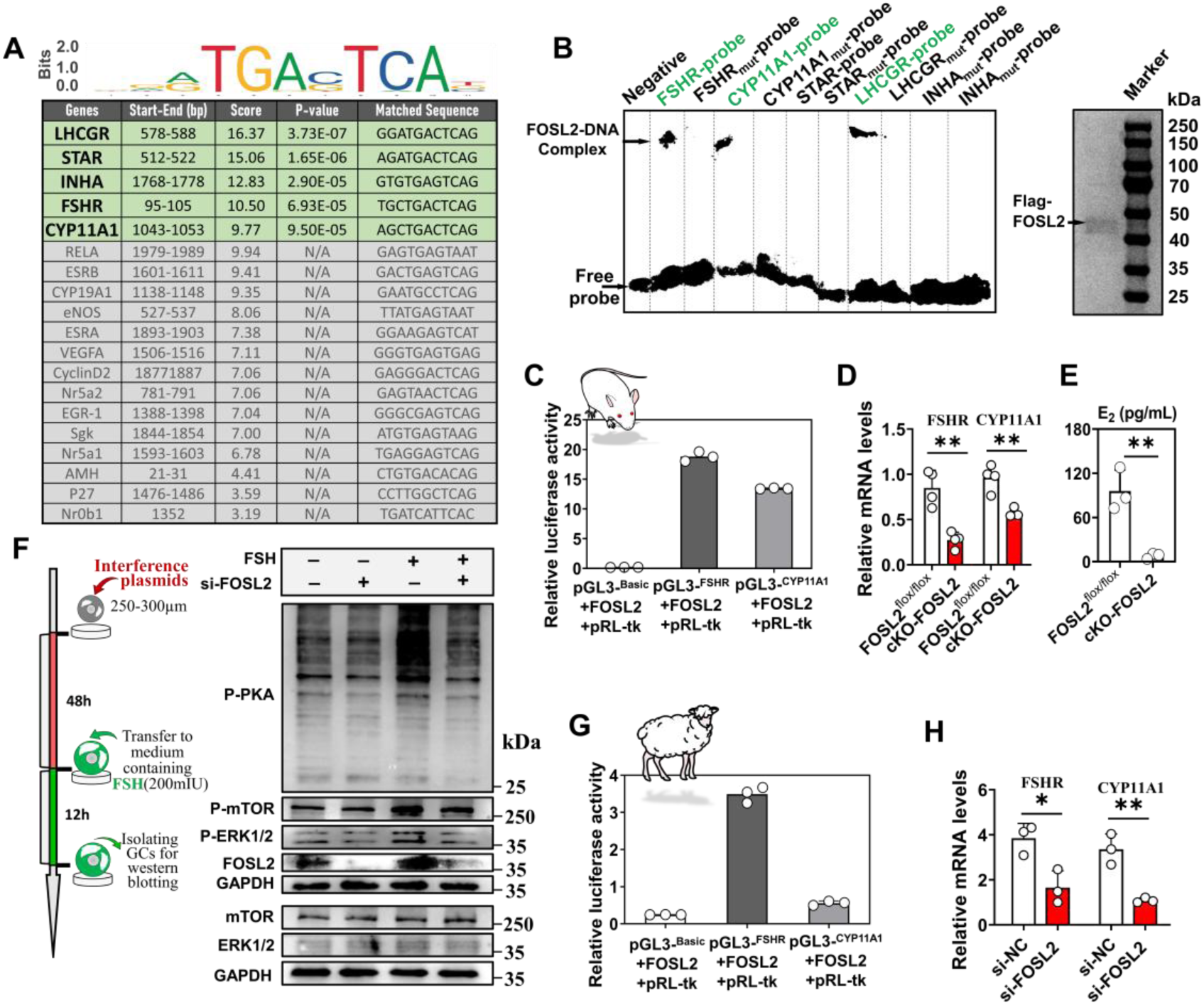
FOSL2 amplified FSH/FSHR signaling through transcriptionally reinforcing the expression of FSHR and CYP11A1. (A) Bioinformatic prediction of FOSL2 target genes. (B) Validation of the binding between FOSL2 and the promoter regions of predicted target genes using EMSA. Right: purification of Flag-tagged FOSL2 protein. (C) Dual-luciferase reporter assay demonstrating the transcriptional activation of FSHR and CYP11A1 promoters by FOSL2. Assay performed in HEK293T cells; n = 3 independent cell samples. (D) Analysis of FSHR and CYP11A1 expression levels following FOSL2 knockout. GCs were isolated from ovaries collected 48 hours after PMSG injection; n = 4 independent GC samples. (E) Measurement of serum estradiol levels following FOSL2 knockout; n = 3 independent serum samples collected 48 hours post-PMSG injection. (F) Effect of FOSL2 knockdown on canonical FSH downstream signaling pathways. Left: schematic representation illustrating the experimental design. Right: Western blotting analysis of FSH-downstream signaling cascades in si-FOSL2 GCs following FSH supplement. Original blots were provided in Figure S6. (G) Dual-luciferase reporter assay validating the transcriptional activation of FSHR and CYP11A1 promoters by FOSL2 in cultured sheep GCs, n=3 independent GC samples. (H) Expression changes of FSHR and CYP11A1 following FOSL2 knockdown in sheep GCs isolated from small antral follicles; n = 3 independent GC samples. Statistical significance was determined using two-tailed unpaired Student’s t test, values were mean ± SD. Significant differences were denoted by*P<0.05, **P<0.01, ***P<0.005, ****P<0.001. Experiments shown in panels B and F were independently repeated three times, and those in panels C, G, and H were repeated twice, all yielding consistent results.

We performed similar assays in cultured sheep GCs. Dual-luciferase reporter assays confirmed FOSL2-driven transcriptional activity of FSHR and CYP11A1 promoters (Figure 5G). Furthermore, qRT-PCR analysis showed significantly reduced expression of both genes in si-FOSL2 GCs compared to si-NC controls (Figure 5H). Together, these results uncover a conserved regulatory role for FOSL2 in enhancing FSH/FSHR signaling across species, primarily through the transcriptional upregulation of FSHR and CYP11A1, thereby supporting GTH-dependent folliculogenesis.

### 6. FSH/FSHR signaling activated FOSL2 transcription via the cAMP-PKA-CREB

To elucidate the mechanism by which FSH/FSHR activates FOSL2 transcription, we employed JASPAR database to predict potential TFs binding to the promoters of FOSL2. As shown in Figure 6A, among the top 10 predicted TFs with the highest binding affinity scores, CREB—a key downstream TF of the PKA kinase cascade— was prominently identified. Given that FSHR is a G protein-coupled receptor capable of activating target gene transcription through the PKA kinase cascade, we hypothesized that the cAMP-PKA-CREB cascade mediates FOSL2 transcription. To test this, we conducted experiments using a follicle culture system (Figure 6B). Treatment with forskolin, an adenylate cyclase activator, led to significant increase of P-PKA and FOSL2 in the absence of exogenous FSH. In contrast, introduce of H89, a PKA inhibitor, effectively blocked FSH-induced increases in FOSL2 levels, confirming the involvement of PKA activity in this regulatory process (Figure 6C). To further validate the role of CREB in regulating FOSL2 transcription, we performed CREB knockdown in cultured follicles (Figure 6D). Compared to the controls, CREB knockdown significantly reduced FOSL2 levels (Figure 6E). Lastly, a dual-luciferase reporter assay demonstrated that CREB markedly enhances the transcriptional activity of the FOSL2 promoter (Figure 6F), highlighting its essential role as a transcriptional activator of FOSL2. Altogether, these findings confirm our speculation that FSH/FSHR signaling activates FOSL2 transcription via the cAMP-PKA-CREB cascade.

**Figure 6.**
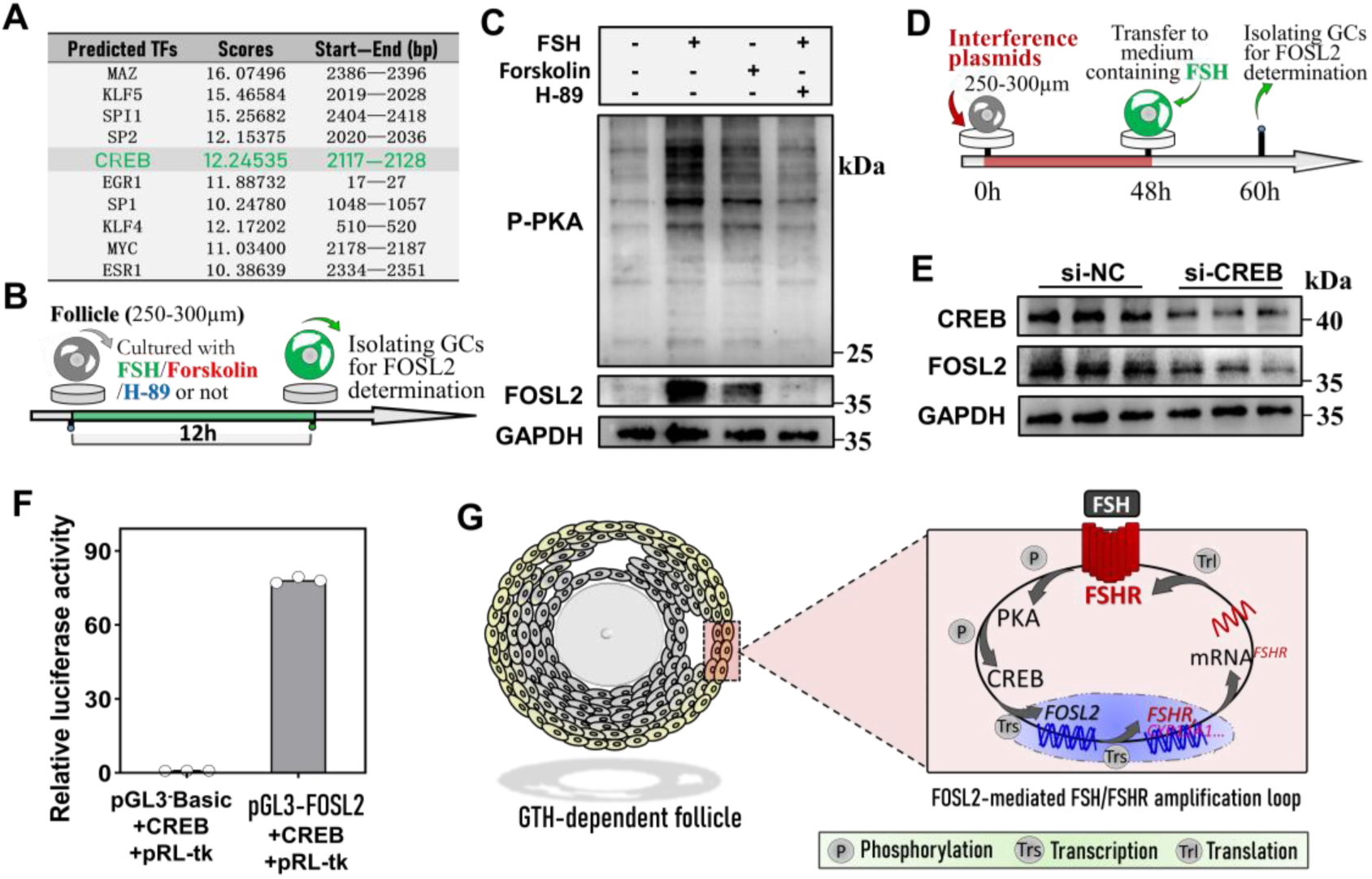
FSH/FSHR signaling activated FOSL2 transcription via the cAMP-PKA-CREB. (A) Prediction of TFs binding to FOSL2’s promoter using JASPAR. (B) Schematic representation illustrating the experimental design for panel C. Forskolin: 20 μM; H-89: 50 μM. (C) Changes in FOSL2 protein contents in GCs following activation or inhibition of the cAMP-PKA cascade. Original blots were provided in Figure S6. (D) Schematic representation illustrating the experimental design for panel E. (E) Changes of FOSL2 protein contents in GCs following CREB knockdown. Original blots were provided in Figure S6. (F) Dual-luciferase reporter assay confirming CREB-mediated transactivation of the FOSL2 promoter. Experiments conducted in HEK293T cells, n=3 independent cell samples. (G) A diagram depicting the FOSL2-mediated amplification loop of FSH/FSHR signaling. During GTH-dependent folliculogenesis, FSH induces FOSL2 expression via the cAMP/PKA/CREB cascade. FOSL2 subsequently upregulates FSHR transcription, establishing an amplification loop that augment FSH/FSHR signaling. Statistical significance was determined using two-tailed unpaired Student’s t test, values were mean ± SD. Significant differences were denoted by *P<0.05, **P<0.01, ***P<0.005, ****P<0.001. The experiments in panels C, E were repeated independently three times, and F was repeated independently twice with consistent results.

## DISCUSSION

FSH/FSHR signaling orchestrates folliculogenesis, with its amplification during GTH-dependent phase being essential for successful follicle maturation. While the mechanisms underlying FSH/FSHR amplification remain incompletely understood, emerging evidence highlights a complex, multilayered regulatory network. Systemically, activin and myostatin enhance FSH secretion, thereby potentiating FSH/FSHR signaling [23,24]. Locally, signaling cascades activated by FSH/FSHR in GCs, such as estrogen signaling, TGF-β/Smad, and Wnt/β-catenin, increase FSHR expression [21,25,26], reinforcing FSH/FSHR signaling. Additionally, GC-derived factors—including insulin-like growth factor 1 (IGF1), nerve growth factor (NGF), and bone morphogenetic proteins (BMPs)—upregulate FSHR expression [27–29], further augmenting FSH/FSHR signaling.

This study identifies FOSL2 as a previously unrecognized regulator critical for FSH/FSHR amplification. As an FSH-target gene, FOSL2 exhibits a spatiotemporal expression pattern parallel to FSHR during the GTH-dependent phase. Through both in vivo and in vitro analyses, we demonstrate that FSH induces FOSL2 expression via the cAMP/PKA/CREB cascade. FOSL2 in turn enhances FSHR transcription through direct promoter binding, establishing a self-amplifying loop. This loop represents a molecular switch, modeled to the all-or-nothing dynamics of GTH-dependent folliculogenesis (Figure 6G). The evolutionary conservation of this mechanism was confirmed through cross-species analyses in sheep, where FOSL2 deficiency similarly attenuated FSH/FSHR signaling and inhibited follicular growth. Genetic ablation of FOSL2 disrupts this loop, halting folliculogenesis at the small antral stage and leading to complete infertility. Interestingly, although FOSL2 knockout mice are infertile, they are still able to ovulate mature oocytes upon superovulation stimulation, with the number reaching nearly 40% of that observed in control mice (Figure S4). This observation suggests that supraphysiological FSH administration may partially compensate for the attenuated FSH/FSHR signaling caused by FOSL2 deficiency. Additionally, FOSL2 also activates the transcription of CYP11A1, a key enzyme involved in steroidogenesis. Disruption of FOSL2 at follicular and organismal levels uniformly impaired estrogen biosynthesis (Figure 5E, S5). Given estrogen’s established capacity to potentiate FSH/FSHR signaling [21], we propose that FOSL2 amplify FSH/FSHR through two complementary mechanisms: direct transcriptional regulation of FSHR and indirect potentiation via CYP11A1-mediated estrogen production.

FOSL2 is known to be a stimulus-responsive TF, activated by a variety of hormonal, growth factor, cytokine, and extracellular matrix signals [30,31]. Early knockout models in mice revealed its essential roles in survival, with global deletion leading to neonatal lethality [32]. Subsequent tissue-specific knockout models have delineated its regulatory roles in osteogenic differentiation [33,34], lipid metabolism [35], pulmonary functional maintenance [36–38], and immune homeostasis [39,40]. Particularly, FOSL2 has garnered significant attention in oncology, where it has been shown to promote tumor progression by regulating proliferation, epithelial-mesenchymal transition, metastasis, therapeutic resistance, tumor microenvironment remodeling, and immune evasion [41–44]. Surprisingly, despite being identified as an GTH-responsive ovarian gene decades ago [45,46], FOSL2’s specific reproductive functions had not been characterized until this study. Our investigation reveals that FOSL2 amplifies FSH/FSHR signaling by upregulating FSHR expression and enhancing estrogen synthesis, thereby specifically regulating GTH-dependent folliculogenesis. This finding fills a critical gap in our understanding of FOSL2’s role in reproduction. Moreover, we propose that FOSL2’s functional scope extends beyond folliculogenesis, encompassing broader aspects of reproductive processes. For instance, EMSA demonstrated LHCGR is another direct target gene of FOSL2 (Figure 5B). This suggest FOSL2 may play a dual regulatory role where it not only transcriptionally activates FSHR to facilitate the growth of GTH-dependent follicles, but also confers ovulatory competence to these follicles via transcriptionally activating LHCGR. Furthermore, the upregulation of FOSL2 in pre-ovulatory follicles in response to the LH surge [47] may indicate its involvement in ovulation or luteinization. In addition, given FOSL2’s regulatory influence on female GCs, we hypothesize that it also plays a significant role in male Sertoli cells, potentially influencing spermatogenic efficiency. These findings open avenues for further investigation into FOSL2’s pleiotropic roles in both reproductive physiology and pathophysiology.

The full spectrum of FOSL2 target genes in GCs likely extends beyond FSHR, CYP11A1 and LHCGR. While we initially attempted to comprehensively identify FOSL2 target genes using ChIP-seq, technical limitations due to suboptimal commercial FOSL2 antibodies led us to employ alternative methodologies. The experimental approach in this study combined EMSA with luciferase reporter assays for targeted verification within a limited candidate gene spectrum, which may have resulted in an underestimation of potential targets. This limitation underscores the need for high-quality, ChIP-grade FOSL2 antibodies to enable comprehensive identification of FOSL2’s target genes, which will be crucial for a more complete understanding of the mechanisms through which FOSL2 regulates GTH-dependent folliculogenesis. According to the classic “two-cell, two-gonadotropin” theory [48], LH stimulates theca cells to express CYP11A1, which converts cholesterol to androgens and transfers them to GCs. In parallel, FSH induces GCs to express CYP19A1, which converts androgens into estrogens. Our findings indicate that FOSL2 directly regulates CYP11A1 transcription in GCs, rather than CYP19A1. Furthermore, specific deletion of FOSL2 in GCs impairs estrogen synthesis. This observation challenges conventional understanding since CYP19A1-mediated aromatization represents the crucial step in estrogen biosynthesis. Three non-exclusive hypotheses warrant further exploration: (i) FOSL2-dependent FSHR upregulation amplifies FSH/FSHR signaling, indirectly promoting estrogen production; (ii) GCs may possess the capacity for de novo androgen synthesis via CYP11A1, akin to theca cell function; or (iii) FOSL2 may regulate additional, yet unidentified genes of the estrogen biosynthetic machinery. These hypotheses await experimental validation, particularly through functional studies of CYP11A1 in GC steroidogenesis and improved ChIP-seq methodologies.

In summary, this study introduces a novel FOSL2-centered amplification loop for FSH/FSHR signaling, emphasizing FOSL2’s indispensable role in female reproductive processes. This loop not only advances the theoretical framework for understanding GTH-dependent folliculogenesis, but also provides valuable insights into the pathophysiology of related reproductive disorders and potential therapeutic avenues.

## MATERIALS AND METHODS

### Experimental design

This study commenced with the prediction of TFs potentially involved in the amplification FSH/FSHR signaling. By integrating single-cell and spatial transcriptomic analyses, candidate TFs exhibiting upregulation in GCs during folliculogenesis were identified. Subsequently, qRT-PCR was employed to screen for FSH-responsive TFs among the candidates. From these, TFs demonstrating a spatiotemporal expression pattern consistent with FSHR during the GTH-dependent phase were selected for further investigation.

Following the identification of FOSL2 as the research subject, knockdown experiments were conducted in GCs derived from both mouse and sheep to examine its role in regulating proliferation, apoptosis, and estrogen synthesis. To investigate the functional relevance of FOSL2 in a more physiologically relevant background, FOSL2 was knocked down in cultured mouse and sheep follicles to determine whether its knockdown selectively disrupts GTH-dependent folliculogenesis and estrogen secretion. In parallel, GC-specific FOSL2 knockout mice were generated to evaluate the in vivo effects of FOSL2 deficiency on GTH-dependent folliculogenesis, estrous cyclicity, estrogen production, and reproductive capacity.

To explore the potential mechanism through which FOSL2 modulates GTH-dependent folliculogenesis—specifically its role in amplifying FSH/FSHR signaling— bioinformatic prediction was performed to evaluate FOSL2’s binding affinity to the FSHR promoter. This prediction was experimentally validated using EMSA and luciferase reporter assays. Furthermore, FOSL2 knockdown was carried out in cultured GTH-dependent follicles, followed by Western blotting to assess the resulting changes in the activity of three downstream signaling pathways of FSH/FSHR: cAMP-PKA, mTOR, RAS-RAF-ERK.

Finally, a combination of Western blotting, RNA interference, and luciferase reporter assays was employed in a follicle culture system to elucidate the downstream signaling cascade through which FSH/FSHR upregulates FOSL2 expression.

### Animals

Kunming mice were purchased from the Center for Animal Testing of Huazhong Agricultural University (Wuhan, China). FOSL2^flox/flox^ C57BL/6J mice were purchased from GemPharmatech Co., Ltd. (Nanjing, China), and FSHR-Cre mice were donated by Dr. Y.S. (Shandong University, China). Following the mating of FOSL2*^flox/flox^* mice with FSHR-Cre mice, specific deletion of exons 2 to 4 of FOSL2 in GCs can be achieved. Mice were reared in an SPF laboratory animal house, at a constant temperature of 22±2 ℃, being allowed to access food and water ad libitum with 12h light-dark cycles. Sheep ovaries were purchased from slaughterhouses. Prior approval from the Institutional Animal Ethics Committee of Huazhong Agricultural University was obtained, with the approved protocol number being HZAUMO-2022-0181 (mouse); HZAUSH-2025-0002 (sheep).

### Estrus cycle determination

Vaginal smears were performed to determine the estrous cycle stage in mice. Briefly, 50 μL of physiological saline was introduced into the vaginal canal and gently aspirated three times to obtain cellular samples. The lavage fluid was then transferred onto glass slides and air-dried. The samples were stained by H&E staining kit (Servicebio, China). After washing with distilled water, the slides were air-dried completely and examined under a light microscope. The estrous cycle stage (proestrus, estrus, metestrus, or diestrus) was determined based on the predominant cell types observed: nucleated epithelial cells, cornified epithelial cells, leukocytes, or their combinations [49].

### Superovulation

To induce GTH-dependent folliculogenesis, female mice were intraperitoneally injected with 5 IU of PMSG (Ningbo Sansheng Biological Technology, China). After 48 hours, ovaries were either harvested for morphological assessment or the mice received an intraperitoneal injection of 5 IU human chorionic gonadotropin (hCG, Ningbo Sansheng Biological Technology, China) to trigger ovulation. For oocyte collection, oviducts were dissected 16 hours post-hCG administration. Cumulus-oocyte complexes were collected from the oviductal ampullae and treated with 0.3% hyaluronidase (Sigma-Aldrich, USA) to disperse cumulus cells. The number of ovulated oocytes was then quantified under stereomicroscopic observation (Nikon, Japan).

### Integrative analysis of scRNA-seq and spatial transcriptomics

While scRNA-seq provides high-resolution transcriptional profiles of GCs, it exhibits limited accuracy in distinguishing GCs across distinct follicular stages. Conversely, spatial transcriptomics precisely resolves follicular staging but lacks the depth of scRNA-seq. To address these limitations, we integrated both modalities to achieve high-resolution transcriptomic mapping of GCs across follicular development. Ovaries from PMSG-injected mice (48 hours post-injection), containing follicles at all developmental stages, were collected for single-cell RNA sequencing (10× Genomics platform) by Yingzi Gene (Wuhan, China). Cell clustering was performed using a graph-based approach, which involved constructing a sparse nearest-neighbor graph followed by Louvain Modularity Optimization for community detection [50]. Differential gene expression analysis between cell clusters was conducted using the Seurat Bimod test, with significance thresholds set at a false discovery rate (FDR) ≤ 0.05 and an absolute log2 fold change ≥ 1.5. Spatial transcriptomic data from mouse ovaries (48 hours post-PMSG injection), originally generated by Mantri et al., were retrieved from the Gene Expression Omnibus (GEO) database (Accession: GSE240271), with processed metadata and cell annotation files obtained from the associated GitHub repository (https://github.com/madhavmantri/mouse_ovulation).

The workflow of integrative analysis began with data preprocessing, where total Unique Molecular Identifier (UMI) counts in scRNA-seq data were normalized using scanpy.pp.normalize_total to eliminate variability in sequencing depth. Spatial mapping was then performed using a deep learning model (*tangram.map_cells_to_space*) implemented in PyTorch, trained on 189 ovarian marker genes [51]. The model underwent 2,000 epochs with a learning rate of 0.01, operating in "CELLS" mode for cell-to-spot mapping, with spatial constraints enforced via *Density_prior=’rna_count_based*’. Optimization was performed using the Adam optimizer to minimize *Kullback-Leibler (KL)* divergence between scRNA-seq and spatial transcriptomic datasets. Cell annotations (Subclass_label) were projected onto spatial coordinates using *tg.project_cell_annotations*, with dominant cell types per spot determined via *tangram_ct_pred*. Integrated transcriptomes were generated using *tg.project_genes*, followed by rigorous quality assessment. Model performance was evaluated using *tg.plot_training_scores*, whil*e tg.plot_genes_sc* was used to compare spatial and integrated transcriptomes to validate low-scoring gene expression patterns. Concordance between scRNA-seq and integrated transcriptomes was further confirmed by calculating gene Area Under the Curve (AUC) scores (*tg.compare_spatial_geneexp*). Following integration, the high-resolution and spatially informed transcriptional atlas of GCs encompassing the developmental continuum from Type-5b, Type-6, and Type-7/8 follicles were extracted. Upregulated genes were identified and cross-referenced with the AnimalTFDB database (http://bioinfo.life.hust.edu.cn/AnimalTFDB/) to determine the upregulated TFs. Follicle staging criteria was established by the Pedersen and Peters. Type-5b follicles contain ≥5 GC layers surrounding a fully grown oocyte without a follicular antrum; Type-6 follicles exhibit multilayered GCs separated by small, irregular antra; Type-7 follicles possess a single large antrum with a defined cumulus oophorus; and Type-8 follicles display a large antrum with a well-developed cumulus stalk [52].

### Cell and follicle culture

Sheep GCs were aseptically isolated from small antral follicles (300-350 μm diameter). Mouse KK1 GC line was donated by Dr. X.C., and X.M. (Chongqing Medical University, China). Cells were cultured in DMEM/F12 medium (Gibco, USA) supplemented with 10% fetal bovine serum (Gibco, USA) and 100 U/mL penicillin/streptomycin (Gibco, USA), and maintained at 37 ℃ in a humidified atmosphere of 5% CO₂.

Mouse Type-5b (150-180 μm diameter) and Type-6 follicles (250-300 μm diameter) were microdissected from mouse ovaries using 33-gauge microneedles (KONSFI, China). Follicles were individually cultured in 96-well plates (BKMAM, China) containing 50 μL of culture medium under mineral oil (Sigma-Aldrich, USA) overlay. The culture medium consisted of α-MEM (Gibco, USA) supplemented with 1% ITS-G (Macklin, China), 5% FBS (Serana, Germany), 10 mIU/mL FSH (NSHF, China), 100 U/mL penicillin/streptomycin (Servicebio, China). The culture system was maintained at 37 ℃ with 5% CO₂. Following 120 hours of culture, the small antra are formed within the Type-5b follicles. After 96 hours of culture, the Type-6 follicles develop to the pre-ovulatory stage.

Small antral follicles (300-350 μm diameter) were isolated from sheep ovaries using ophthalmic scissors and 26-gauge microneedles (KONSFI, China). Follicles were cultured in 96-well plates containing 100 μL of culture medium under mineral oil overlay at 38.5 ℃ with 5% CO₂. The culture medium contained α-MEM basal medium, 10% FBS, 1% ITS, 50 μg/mL ascorbic acid (Sigma-Aldrich, USA), 2 mM hypoxanthine (Sigma-Aldrich, USA), 2 mM glutamine (Sigma-Aldrich, USA), 10 mIU/mL FSH, 100 U/mL penicillin/streptomycin. Under these conditions, follicle diameter typically increased to approximately 500 μm after 96 hours of culture.

### RNA Interference

FOSL2-specific siRNA (Genepharma, China) was employed to silence FOSL2 expression in cultured KK1 GC line. Transfection was performed using jetPRIME® transfection reagent (PolyPlus-transfection, France) according to the manufacturer’s protocol. Briefly, GCs were transfected with 100 pmol of siRNA in culture medium for 48 hours, followed by replacement with fresh culture medium for subsequent experiments.

Lentivirus-mediated RNA interference was used to decrease the expression of FOSL2 and CREB in cultured follicles or sheep GCs. The PLKO.1-EGFP-PURO plasmid (Genecreate, China) was used to construct shRNA interference vectors. Lentiviral particles were produced by co-transfecting HEK293T cells (ATCC, USA) with 4.8 μg of interference vector, 2.4 μg of pMD2.G envelope plasmid (Addgene, USA), 3.6 μg of pSPAX2 packaging plasmid (Addgene, USA). Viral supernatants were collected 48 hours post-transfection, centrifuged at 4000r for 10 minutes, and filtered through 0.45 μm PVDF membranes (Sigma-Aldrich, USA). Follicles and sheep GCs were then incubated with the prepared viral particles (titer: 1.25 × 10⁷ particles/mL) in culture medium containing 10 μg/mL Polybrene (Sigma-Aldrich, USA). Transfection durations were optimized as 48 hours (mouse follicles and sheep GCs) and 96 hours (sheep follicles), respectively. Transfection was confirmed by EGFP fluorescence visualization. Following transfection, samples were either immediately processed for phenotypic analysis or maintained in fresh culture medium for subsequent experiments. Non-targeting siRNA were purchased from Sigma-Aldrich (USA). The siRNA target sequences were provided in Table S5.

### H&E Staining

Ovaries were fixed in 4% paraformaldehyde (Servicebio, China) for 24 hours at 4 ℃, followed by standard paraffin embedding. Sections (5 μm thickness) were prepared using a rotary microtome (Leica, Germany). After deparaffinization and rehydration, the sections were stained with hematoxylin (Servicebio, China) for 5 minutes. Then the sections were stained with eosin (Servicebio, China) for 5 minutes and followed by dehydration with graded alcohol and clearing in xylene. Stained sections were examined under a microscope (Olympus, Japan) equipped with a digital camera. Morphometric analysis was performed using ImageJ software (v1.52, NIH, USA).

### qRT-PCR assay

Total RNA from samples was extracted using TRIzol reagent (Takara, Japan). Reverse transcription was performed using the PrimeScript RT reagent kit (Takara, Japan). The qRT-PCR was performed by CFX384 Real-Time PCR System (Bio-Rad, USA). Reaction system consist of 5 μL SYBR Green (Biosharp, China), 2 μL complementary DNA template, 250 nM of the forward and reverse primers for each, and ddH_2_O was supplemented to a total volume of 10 μL. The reaction protocol was conducted as described: an initial denaturation step at 95 ℃ for 10 minutes, succeeded by 35 cycles comprising denaturation at 95 ℃ for 10 s and annealing/ extension at 60 ℃ for 30 seconds. A final step included a melting curve analysis ranging from 60 ℃ to 95 ℃, with a 0.5 ℃ increment every 5 seconds. Using ACTB to normalize gene expression levels, and using comparative 2^−ΔΔCt^ method to determine relative RNA. The primer sequences used for PCR amplification are provided in Table S5.

### Western blotting

Total protein was extracted using RIPA lysis buffer (ComWin Biotech, China) supplemented with protease and phosphatase inhibitor (ComWin Biotech, China). Protein concentrations were quantified using the BCA Protein Assay Kit (Servicebio, China) according to the manufacturer’s protocol. Equal amounts of protein (typically 20-50 μg per lane) were separated by 10% SDS-PAGE and subsequently transferred to polyvinylidene fluoride (PVDF) membranes (0.45 μm pore size; Sigma-Aldrich, USA) using standard wet transfer methods. Membranes were blocked with 5% non-fat dry milk in Tris-buffered saline containing 0.1% Tween-20 (TBST) for 2 hours at room temperature. Following blocking, membranes were incubated overnight at 4°C with the following primary antibodies diluted in blocking buffer: FOSL2 (1:100, Sigma-Aldrich, USA), Caspase-3 (1:100, CST, USA), Cleaved-Caspase-3 (1:100, CST, USA), PARP (1:100, CST, USA), Cleaved-PARP (1:100, CST, USA), Bax (1:100, CST, USA), phospho-PKA (1:100, CST, USA), mTOR (1:100, CST, USA), phospho-mTOR (1:100, CST, USA), ERK1/2 (1:100, Servicebio, China), phospho-ERK1/2 (1:100, Abclonal, China), CREB (1:100, CST, USA), phospho-CREB (1:100, CST, USA), α-Tubulin (1:100, Servicebio, China), β-Actin (1:100, Abclonal, China), and GAPDH (1:100, Abclonal, China). After primary antibody incubation, membranes were washed three times (10 min each) with TBST and then incubated with horseradish peroxidase (HRP)-conjugated secondary antibodies (goat anti-rabbit IgG, 1:400; goat anti-mouse IgG, 1:400, Biodragon-immunotech, China) for 1 hour at room temperature. Protein bands were visualized using an enhanced chemiluminescence (ECL) detection kit (Biosharp, China) according to the manufacturer’s instructions, and images were captured using a ChemiDoc imaging system (Tanon-520, China). Band intensities were quantified using ImageJ software (NIH, USA), with normalization to appropriate loading controls (α-Tubulin, β-Actin, or GAPDH).

### Immunofluorescence

Ovarian tissues were sectioned at 5 μm thickness using a rotary microtome and mounted on poly-L-lysine-coated slides (CITOTEST, China). After deparaffinization through xylene and graded ethanol series, antigen retrieval was performed in citrate buffer at 95-98 ℃ for 25 minutes. Sections were permeabilized with 0.5% Triton X-100 in PBS for 20 minutes at room temperature, followed by blocking with 10% goat serum (Boster, China) in PBS for 60 minutes at room temperature. For immunostaining, sections were incubated with rabbit anti-FOSL2 primary antibody (1:200 dilution, Sigma-Aldrich, USA) at 4°C overnight, then with an HRP-conjugated secondary antibody working solution (Recordbio Biological Technology, China) at 37 ℃ for 50 minutes. Subsequently, the sections were treated with the fluorescent dye TYR-520 (Recordbio Biological Technology, China) for 5 min, followed by a heat treatment to eliminate nonspecific dye binding. Dual-staining of FOSL2 and FOXL2 was conducted using the Three-color Fluorescence kit (Recordbio Biological Technology, China) based on Tyramide Signal Amplification technology. Briefly, following antigen blocking, the sections were incubated overnight at 4 ℃ with rabbit anti-FOSL2 primary antibody. After rinsing with PBS, the sections were incubated with an HRP-conjugated secondary antibody working solution (Recordbio Biological Technology, China) at 37 ℃ for 50 minutes. Subsequently, the sections were treated with the fluorescent dye TYR-520 for 5 min, followed by eliminating nonspecific dye binding. The same protocol was applied for FOXL2 detection using the primary antibody against FOXL2 (1:100 dilution, Abcam, USA) and fluorescent dye TYR-570 (Recordbio Biological Technology, China) for FOXL2 staining. Additionally, cell nuclei were stained with DAPI (Biosharp, China). Following another round of washing, the sections were imaged using an LSM800 confocal microscope system (Zeiss, Germany) and the resulting images were analyzed using Zen 2.3 lite software.

### EdU Staining

GC proliferation was evaluated using the EdU assay kit (RiboBio, China). Following siRNA-mediated gene silencing, GCs grown on cell climbing slices (RiboBio, China) were labeled with 50 μM EdU for 2 hours at 37 ℃ under 5% CO₂ atmosphere. Subsequently, GCs were fixed with 4% paraformaldehyde (Servicebio, China) for 30 minutes at room temperature. Permeabilization was achieved using 0.5% Triton X-100 (Servicebio, China) in PBS for 10 minutes. The reaction was performed by incubating samples with EdU dye reaction solution for 30 minutes in the dark, followed by nuclear counterstaining with Hoechst 33342 (RiboBio, China) for 10 minutes. For whole follicle assessment, cultured follicles were incubated with 50 μM EdU for 24 hours post-plasmid transfection. Follicles were then embedded in optimal cutting temperature compound (OCT) (Sakura Finetek, Japan) and snap-frozen in liquid nitrogen for 1 minute. Cryosections (5 μm thickness) were obtained using a CM195 cryostat (Leica, Germany). The sections were processed for EdU detection using the same reaction conditions as for GCs, with 30-minute EdU staining and 10-minute Hoechst 33342 counterstaining. Fluorescent images were captured using a fluorescence microscope (Olympus, Japan). Quantitative analysis was performed using ImageJ software. Calculation of proliferation index as (EdU^+^ cells/total Hoechst^+^ cells) × 100%.

### TUNEL Staining

Following siRNA transfection, GCs grown on cell climbing slices were fixed with 4% paraformaldehyde for 15 minutes at room temperature and washed three times with PBS. Cell membranes were permeabilized with 0.5 % Triton X-100 in PBS at 37 ℃ for 20 minutes. After PBS washing, samples were equilibrated with 100 μL TdT Equilibration Buffer (Roche, Switzerland) at 37 ℃ for 30 minutes. The terminal deoxynucleotidyl transferase (TdT)-mediated dUTP nick-end labeling reaction was performed by incubating samples with TUNEL reaction mixture (Roche, Switzerland) at 37 ℃ for 60 minutes in a humidified dark chamber. Nuclei were counterstained with DAPI for 10 minutes, and slides were mounted with anti-fade mounting medium. Fluorescence images were captured using a microscope (Olympus, Japan). Apoptotic cells were identified by green fluorescence (FITC signal). The apoptosis rate was calculated as: (number of TUNEL^+^ cells / total number of DAPI-stained cells) × 100%.

### Flow Cytometry Assay

The cell cycle distribution was assessed using a Cell Cycle Detection Kit (Keygentec, China). Following RNA interference, GCs were fixed in 70% ice-cold ethanol at 4 ℃ overnight. Fixed cells were then washed with PBS and stained with 500 µL propidium iodide (PI)/RNase A working solution for 60 minutes at room temperature in the dark. Cellular DNA content was measured by FACSVerse system (BD Biosciences, USA) with excitation at 488 nm and emission detection at 617 nm. Data analysis was performed using ModFit LT™ software version 4.1 (Verity Software House, USA) to determine the percentage of cells in G1, S, and G2 phases.

Apoptotic cells were quantified using an Annexin V-FITC/PI apoptosis detection kit (Beyotime, China). Briefly, GCs were resuspended in 195 µL Annexin V binding buffer and stained with 5 µL Annexin V-FITC and 10 µL PI working solution for 15 minutes at room temperature in the dark. Flow cytometric analysis was immediately performed using the FACSVerse system, with Annexin V-FITC fluorescence detected in the FL1 channel (530/30 nm bandpass filter) and PI fluorescence in the FL3 channel (>670 nm longpass filter). Viable (Annexin V^-^/PI^-^), early apoptotic (Annexin V^+^/PI^-^), late apoptotic (Annexin V^+^/PI^+^), and necrotic (Annexin V^-^/PI^+^) cell populations were distinguished and quantified using FlowJo software (Leonard Herzenberg, USA).

### RTCA assay

Cellular proliferation dynamics were quantitatively assessed using the xCELLigence RTCA DP Instrument (Roche, Switzerland), which employs impedance-based detection to monitor cell proliferation in real time. Briefly, KK1 GCs were seeded into E-plate 16 (ACEA Biosciences, USA) at an optimized density of 1×10^5^ cells/mL, with 100 μL of cell suspension dispensed into each well. After a 30-minute incubation at room temperature to ensure proper cell sedimentation and adherence, the E-plate 16 was placed in the RTCA station for continuous impedance monitoring. The system automatically recorded impedance changes, expressed as a dimensionless cell index value, with measurements acquired every 15 minutes throughout the 160-hour experimental duration. To evaluate the impact of FOSL2 knockdown on proliferation kinetics, si-RNA transfection was performed at the 20-hour time point following monitoring initiation.

### EMSA Assay

The coding sequence of FOSL2 was subcloned into the pcDNA3.1-3XFlag expression vector (Addgene, USA) to generate Flag-tagged FOSL2 protein. Following overexpression, Flag-FOSL2 protein were immunoprecipitated using anti-Flag monoclonal antibody (Beyotime, China) and subsequently eluted under acidic conditions (0.1 M glycine, pH 2.7), followed by immediate neutralization with 1 M Tris-HCl (pH 8.5). The DNA EMSA was performed using the Chemiluminescent EMSA Kit (Beyotime, China), according to the manufacturer’s instructions. Briefly, biotin-labeled double-stranded DNA probes (Genecreate, China) were incubated with purified Flag-FOSL2 protein in binding buffer for 30 minutes at room temperature to allow complex formation. Protein-DNA interactions were resolved on a 4% non-denaturing polyacrylamide gel at 100 V for 60 minutes. The resolved complexes were then electrophoretically transferred to Amersham Hybond-N^+^ nylon membranes (Cytiva, USA) and detected using HRP-conjugated streptavidin followed by autoradiography. The specific primer sequences used for probe generation were provided in Table S5.

### Luciferase Reporter Gene Assay

The promoter regions of FSHR, CYP11A1 and FOSL2 were PCR-amplified and directionally cloned into the pGL3-Basic luciferase reporter vector (Promega, USA) using the ClonExpress Ultra One Step Cloning Kit (Vazyme, China). For over-expression vector construction, the complete coding sequences of FOSL2 and CREB were amplified and subcloned into pcDNA3.1-3XFlag vectors (Addgene, USA). HEK293T cells were plated in 24-well plates and cultured for 24 hours prior to transfection. Cells were co-transfected with 248 ng of over-expression vector, 248 ng of pGL3-based reporter construct, and 2.5 ng of pRL-TK Renilla luciferase control vector (Promega, USA) using jetPRIME transfection reagent according to the manufacturer’s protocol. After 24-hour incubation, cells were harvested in 100 μL passive lysis buffer and analyzed using the Dual-Luciferase Reporter Assay System (Promega, USA). Firefly and Renilla luciferase activities were quantified sequentially using an EnSpire Multimode Plate Reader (PerkinElmer, USA). Relative promoter activity was calculated as the ratio of firefly to Renilla luciferase luminescence. All primer sequences used for cloning and mutagenesis are provided in Table S5.

### Hormone Determination

Serum and culture medium estradiol (E2) concentrations were quantitatively determined using a highly specific radioimmunoassay (RIA) system. All measurements were performed by the Beijing North Institute of Biological Technology (Beijing, China), a certified clinical testing laboratory, utilizing a commercially available RIA kit (Bioengineering Institute, China) with established sensitivity (5 pg/mL). The assay employed competitive binding principles using ^125^I-labeled estradiol tracer and highly specific anti-estradiol antibodies, following the manufacturer’s standardized protocol.

### Statistics Analysis

Statistical analyses were using GraphPad Prism 10.0 (GraphPad). Data were expressed as the mean ± SD. Two-tailed unpaired Student’s t test and one-way analysis of variance followed by Tukey’s post hoc test were used to analyze the statistical significance between two groups and among multiple groups, respectively. Chi-squared test was used in the comparison between the percentages. The statistical significance was set at P-value <0.05.

## DATA AVAILABILITY

The RNA-Seq data reported in this study have been deposited in the Gene Expression Omnibus database. All data are available from the corresponding author upon reasonable request.

## SUPPORTING INFORMATION

This article contains supporting information.

## ACKNOWLEDGEMENTS

We are grateful to Prof. Dr. Louis Dubeau (University of South California, USA) for donating the FSHR-Cre mice, Dr. X.C., and X.M. (Chongqing Medical University, China) Dr. X.C. for donating the KK1 GC line.

## AUTHORS’ CONTRIBUTION

C.H. conceived, designed, conducted the experiments, analyzed and interpreted the data; H.S., C.C. and Z.R. anticipated in experiment design and conduction, data analysis, and manuscript preparation; J.L., Z.W., X.W., Y.Z., W.K., B.T. and Y.L. assisted with sample collection and experiments conduction; Y.S. bred FSHR-Cre mice; C.H., H.S., W.R. C.C. and Z.R. wrote the manuscript; X.L. and W. R. improved the manuscript. C.H., X.L., Y.S. funded this project. C.H., X.L., W.R. supervised this project. All authors approved the final version.

## REFERENCES

1. Njagi, P., Groot, W., Arsenijevic, J., Dyer, S., Mburu, G., & Kiarie, J. (2023). Financial costs of assisted reproductive technology for patients in low- and middle-income countries: a systematic review. Human reproduction open, 2023(2), hoad007. 10.1093/hropen/hoad007.

2. Wang, X., Zhou, S., Wu, Z., Liu, R., Ran, Z., Liao, J., Shi, H., Wang, F., Chen, J., Liu, G., et al. (2023). The FSH-mTOR-CNP signaling axis initiates follicular antrum formation by regulating tight junction, ion pumps, and aquaporins. The Journal of biological chemistry, 299(8), 105015. 10.1016/j.jbc.2023.105015.

3. Aerts, J. M., & Bols, P. E. (2010). Ovarian follicular dynamics: a review with emphasis on the bovine species. Part I: Folliculogenesis and pre-antral follicle development. Reproduction in domestic animals = Zuchthygiene, 45(1), 171–179. 10.1111/j.1439-0531.2008.01302.x.

4. Aerts, J. M., & Bols, P. E. (2010). Ovarian follicular dynamics. A review with emphasis on the bovine species. Part II: Antral development, exogenous influence and future prospects. Reproduction in domestic animals = Zuchthygiene, 45(1), 180–187. 10.1111/j.1439-0531.2008.01298.x.

5. Dewailly, D., Robin, G., Peigne, M., Decanter, C., Pigny, P., & Catteau-Jonard, S. (2016). Interactions between androgens, FSH, anti-Müllerian hormone and estradiol during folliculogenesis in the human normal and polycystic ovary. Human reproduction update, 22(6), 709–724. 10.1093/humupd/dmw027.

6. Ulloa-Aguirre, A., Reiter, E., & Crépieux, P. (2018). FSH Receptor Signaling: Complexity of Interactions and Signal Diversity. Endocrinology, 159(8), 3020–3035. 10.1210/en.2018-00452.

7. Hunzicker-Dunn, M., & Maizels, E. T. (2006). FSH signaling pathways in immature granulosa cells that regulate target gene expression: branching out from protein kinase A. Cellular signalling, 18(9), 1351–1359. 10.1016/j.cellsig.2006.02.011.

8. Lou, L., Urbani, J., Ribeiro-Neto, F., & Altschuler, D. L. (2002). cAMP inhibition of Akt is mediated by activated and phosphorylated Rap1b. The Journal of biological chemistry, 277(36), 32799–32806. 10.1074/jbc.M201491200

9. Alam, H., Maizels, E. T., Park, Y., Ghaey, S., Feiger, Z. J., Chandel, N. S., & Hunzicker-Dunn, M. (2004). Follicle-stimulating hormone activation of hypoxia-inducible factor-1 by the phosphatidylinositol 3-kinase/AKT/Ras homolog enriched in brain (Rheb)/mammalian target of rapamycin (mTOR) pathway is necessary for induction of select protein markers of follicular differentiation. The Journal of biological chemistry, 279(19), 19431–19440. 10.1074/jbc.M401235200.

10. Liu, L., Hao, M., Zhang, J., Chen, Z., Zhou, J., Wang, C., Zhang, H., & Wang, J. (2023). FSHR-mTOR-HIF1 signaling alleviates mouse follicles from AMPK-induced atresia. Cell reports, 42(10), 113158. 10.1016/j.celrep.2023.113158.

11. Gonzalez-Robayna, I. J., Falender, A. E., Ochsner, S., Firestone, G. L., & Richards, J. S. (2000). Follicle-Stimulating hormone (FSH) stimulates phosphorylation and activation of protein kinase B (PKB/Akt) and serum and glucocorticoid-lnduced kinase (Sgk): evidence for A kinase-independent signaling by FSH in granulosa cells. Molecular endocrinology (Baltimore, Md.), 14(8), 1283–1300. 10.1210/mend.14.8.0500.

12. Park, Y., Maizels, E. T., Feiger, Z. J., Alam, H., Peters, C. A., Woodruff, T. K., Unterman, T. G., Lee, E. J., Jameson, J. L., & Hunzicker-Dunn, M. (2005). Induction of cyclin D2 in rat granulosa cells requires FSH-dependent relief from FOXO1 repression coupled with positive signals from Smad. The Journal of biological chemistry, 280(10), 9135–9148. 10.1074/jbc.M409486200.

13. Wang, X. L., Wu, Y., Tan, L. B., Tian, Z., Liu, J. H., Zhu, D. S., & Zeng, S. M. (2012). Follicle-stimulating hormone regulates pro-apoptotic protein Bcl-2-interacting mediator of cell death-extra long (BimEL)-induced porcine granulosa cell apoptosis. The Journal of biological chemistry, 287(13), 10166–10177. 10.1074/jbc.M111.293274.

14. Cottom, J., Salvador, L. M., Maizels, E. T., Reierstad, S., Park, Y., Carr, D. W., Davare, M. A., Hell, J. W., Palmer, S. S., Dent, P., Kawakatsu, H., Ogata, M., & Hunzicker-Dunn, M. (2003). Follicle-stimulating hormone activates extracellular signal-regulated kinase but not extracellular signal-regulated kinase kinase through a 100-kDa phosphotyrosine phosphatase. The Journal of biological chemistry, 278(9), 7167–7179. 10.1074/jbc.M203901200.

15. Hsieh, M., Johnson, M. A., Greenberg, N. M., & Richards, J. S. (2002). Regulated expression of Wnts and Frizzleds at specific stages of follicular development in the rodent ovary. Endocrinology, 143(3), 898–908. 10.1210/endo.143.3.8684.

16. Wang, Y., Chan, S., & Tsang, B. K. (2002). Involvement of inhibitory nuclear factor-kappaB (NFkappaB)-independent NFkappaB activation in the gonadotropic regulation of X-linked inhibitor of apoptosis expression during ovarian follicular development in vitro. Endocrinology, 143(7), 2732–2740. 10.1210/endo.143.7.8902.

17. Jayes, F. C., Day, R. N., Garmey, J. C., Urban, R. J., Zhang, G., & Veldhuis, J. D. (2000). Calcium ions positively modulate follicle-stimulating hormone- and exogenous cyclic 3’,5’-adenosine monophosphate-driven transcription of the P450(scc) gene in porcine granulosa cells. Endocrinology, 141(7), 2377–2384. 10.1210/endo.141.7.7558.

18. Chen, Y., Wang, X., Yang, C., Liu, Q., Ran, Z., Li, X., & He, C. (2020). A mouse model reveals the events and underlying regulatory signals during the gonadotrophin-dependent phase of follicle development. Molecular human reproduction, 26(12), 920–937. 10.1093/molehr/gaaa069.

19. Walters, K., Baldwin, A., Liu, Z., Larsen, M., Mukherjee, N., & Kumar, T. R. (2025). Identification of FSH-regulated and estrous stage-specific transcriptional networks in mouse ovaries. Proceedings of the National Academy of Sciences of the United States of America, 122(7), e2411977122. 10.1073/pnas.2411977122.

20. Chen, H., & Chan, H. C. (2017). Amplification of FSH signalling by CFTR and nuclear soluble adenylyl cyclase in the ovary. Clinical and experimental pharmacology & physiology, 44 Suppl 1, 78–85. 10.1111/1440-1681.12756.

21. Otsuka, F., Moore, R. K., Wang, X., Sharma, S., Miyoshi, T., & Shimasaki, S. (2005). Essential role of the oocyte in estrogen amplification of follicle-stimulating hormone signaling in granulosa cells. Endocrinology, 146(8), 3362–3367. 10.1210/en.2005-0349.

22. Mantri, M., Zhang, H. H., Spanos, E., Ren, Y. A., & De Vlaminck, I. (2024). A spatiotemporal molecular atlas of the ovulating mouse ovary. Proceedings of the National Academy of Sciences of the United States of America, 121(5), e2317418121. 10.1073/pnas.2317418121.

23. Bloise, E., Ciarmela, P., Dela Cruz, C., Luisi, S., Petraglia, F., & Reis, F. M. (2019). Activin A in Mammalian Physiology. Physiological reviews, 99(1), 739–780. 10.1152/physrev.00002.2018.

24. Ongaro, L., Zhou, X., Wang, Y., Schultz, H., Zhou, Z., Buddle, E. R. S., Brûlé, E., Lin, Y. F., Schang, G., Hagg, A., et al. (2025). Muscle-derived myostatin is a major endocrine driver of follicle-stimulating hormone synthesis. Science (New York, N.Y.), 387(6731), 329–336. 10.1126/science.adi4736.

25. Miyoshi, T., Otsuka, F., Suzuki, J., Takeda, M., Inagaki, K., Kano, Y., Otani, H., Mimura, Y., Ogura, T., & Makino, H. (2006). Mutual regulation of follicle-stimulating hormone signaling and bone morphogenetic protein system in human granulosa cells. Biology of reproduction, 74(6), 1073–1082. 10.1095/biolreprod.105.047969.

26. Gupta, P. S., Folger, J. K., Rajput, S. K., Lv, L., Yao, J., Ireland, J. J., & Smith, G. W. (2014). Regulation and regulatory role of WNT signaling in potentiating FSH action during bovine dominant follicle selection. PloS one, 9(6), e100201. 10.1371/journal.pone.0100201.

27. Mani, A. M., Fenwick, M. A., Cheng, Z., Sharma, M. K., Singh, D., & Wathes, D. C. (2010). IGF1 induces up-regulation of steroidogenic and apoptotic regulatory genes via activation of phosphatidylinositol-dependent kinase/AKT in bovine granulosa cells. Reproduction (Cambridge, England), 139(1), 139–151. 10.1530/REP-09-0050.

28. Salas, C., Julio-Pieper, M., Valladares, M., Pommer, R., Vega, M., Mastronardi, C., Kerr, B., Ojeda, S. R., Lara, H. E., & Romero, C. (2006). Nerve growth factor-dependent activation of trkA receptors in the human ovary results in synthesis of follicle-stimulating hormone receptors and estrogen secretion. The Journal of clinical endocrinology and metabolism, 91(6), 2396–2403. 10.1210/jc.2005-1925.

29. Shi, J., Yoshino, O., Osuga, Y., Nishii, O., Yano, T., & Taketani, Y. (2010). Bone morphogenetic protein 7 (BMP-7) increases the expression of follicle-stimulating hormone (FSH) receptor in human granulosa cells. Fertility and sterility, 93(4), 1273–1279. 10.1016/j.fertnstert.2008.11.014.

30. Nishina, H., Sato, H., Suzuki, T., Sato, M., & Iba, H. (1990). Isolation and characterization of fra-2, an additional member of the fos gene family. Proceedings of the National Academy of Sciences of the United States of America, 87(9), 3619–3623. 10.1073/pnas.87.9.3619.

31. Sonobe, M. H., Yoshida, T., Murakami, M., Kameda, T., & Iba, H. (1995). fra-2 promoter can respond to serum-stimulation through AP-1 complexes. Oncogene, 10(4), 689–696.

32. Bozec, A., Bakiri, L., Hoebertz, A., Eferl, R., Schilling, A. F., Komnenovic, V., Scheuch, H., Priemel, M., Stewart, C. L., Amling, M., & Wagner, E. F. (2008). Osteoclast size is controlled by Fra-2 through LIF/LIF-receptor signalling and hypoxia. Nature, 454(7201), 221–225. 10.1038/nature07019.

33. Karreth, F., Hoebertz, A., Scheuch, H., Eferl, R., & Wagner, E. F. (2004). The AP1 transcription factor Fra2 is required for efficient cartilage development. Development (Cambridge, England), 131(22), 5717–5725. 10.1242/dev.01414.

34. Bozec, A., Bakiri, L., Jimenez, M., Rosen, E. D., Catalá-Lehnen, P., Schinke, T., Schett, G., Amling, M., & Wagner, E. F. (2013). Osteoblast-specific expression of Fra-2/AP-1 controls adiponectin and osteocalcin expression and affects metabolism. Journal of cell science, 126(Pt 23), 5432–5440. 10.1242/jcs.134510.

35. Luther, J., Ubieta, K., Hannemann, N., Jimenez, M., Garcia, M., Zech, C., Schett, G., Wagner, E. F., & Bozec, A. (2014). Fra-2/AP-1 controls adipocyte differentiation and survival by regulating PPARγ and hypoxia. Cell death and differentiation, 21(4), 655–664. 10.1038/cdd.2013.198.

36. Ucero, A. C., Bakiri, L., Roediger, B., Suzuki, M., Jimenez, M., Mandal, P., Braghetta, P., Bonaldo, P., Paz-Ares, L., Fustero-Torre, C., et al. (2019). Fra-2-expressing macrophages promote lung fibrosis in mice. The Journal of clinical investigation, 129(8), 3293–3309. 10.1172/JCI125366.

37. Tsujino, K., Li, J. T., Tsukui, T., Ren, X., Bakiri, L., Wagner, E., & Sheppard, D. (2017). Fra-2 negatively regulates postnatal alveolar septation by modulating myofibroblast function. American journal of physiology. Lung cellular and molecular physiology, 313(5), L878–L888. 10.1152/ajplung.00062.2017.

38. Choi, J., Jang, Y. J., Dabrowska, C., Iich, E., Evans, K. V., Hall, H., Janes, S. M., Simons, B. D., Koo, B. K., Kim, J., et al. (2021). Release of Notch activity coordinated by IL-1β signalling confers differentiation plasticity of airway progenitors via Fosl2 during alveolar regeneration. Nature cell biology, 23(9), 953–966. 10.1038/s41556-021-00742-6.

39. Renoux, F., Stellato, M., Haftmann, C., Vogetseder, A., Huang, R., Subramaniam, A., Becker, M. O., Blyszczuk, P., Becher, B., Distler, J. H. W., et al. (2020). The AP1 Transcription Factor Fosl2 Promotes Systemic Autoimmunity and Inflammation by Repressing Treg Development. Cell reports, 31(13), 107826. 10.1016/j.celrep.2020.107826.

40. Lawson, V. J., Maurice, D., Silk, J. D., Cerundolo, V., & Weston, K. (2009). Aberrant selection and function of invariant NKT cells in the absence of AP-1 transcription factor Fra-2. Journal of immunology (Baltimore, Md.: 1950), 183(4), 2575–2584. 10.4049/jimmunol.0803577.

41. Rampioni Vinciguerra, G. L., Capece, M., Scafetta, G., Rentsch, S., Vecchione, A., Lovat, F., & Croce, C. M. (2024). Role of Fra-2 in cancer. Cell death and differentiation, 31(2), 136–149. 10.1038/s41418-023-01248-4.

42. Sarode, P., Zheng, X., Giotopoulou, G. A., Weigert, A., Kuenne, C., Günther, S., Friedrich, A., Gattenlöhner, S., Stiewe, T., Brüne, B., et al. (2020). Reprogramming of tumor-associated macrophages by targeting β-catenin/FOSL2/ARID5A signaling: A potential treatment of lung cancer. Science advances, 6(23), eaaz6105. 10.1126/sciadv.aaz6105.

43. Liu, S. Q., Cheng, X. X., He, S., Xia, T., Li, Y. Q., Peng, W., Zhou, Y. Q., Xu, Z. H., He, M. S., Liu, Y., et al. (2025). Super-enhancer-driven EFNA1 fuels tumor progression in cervical cancer via the FOSL2-Src/AKT/STAT3 axis. The Journal of clinical investigation, 135(8), e177599. 10.1172/JCI177599.

44. Bakiri, L., Hasenfuss, S. C., Guío-Carrión, A., Thomsen, M. K., Hasselblatt, P., & Wagner, E. F. (2024). Liver cancer development driven by the AP-1/c-Jun∼Fra-2 dimer through c-Myc. Proceedings of the National Academy of Sciences of the United States of America, 121(18), e2404188121. 10.1073/pnas.2404188121.

45. Sharma, S. C., & Richards, J. S. (2000). Regulation of AP1 (Jun/Fos) factor expression and activation in ovarian granulosa cells. Relation of JunD and Fra2 to terminal differentiation. The Journal of biological chemistry, 275(43), 33718–33728. 10.1074/jbc.M003555200.

46. Sher, N., Yivgi-Ohana, N., & Orly, J. (2007). Transcriptional regulation of the cholesterol side chain cleavage cytochrome P450 gene (CYP11A1) revisited: binding of GATA, cyclic adenosine 3’,5’-monophosphate response element-binding protein and activating protein (AP)-1 proteins to a distal novel cluster of cis-regulatory elements potentiates AP-2 and steroidogenic factor-1-dependent gene expression in the rodent placenta and ovary. Molecular endocrinology (Baltimore, Md.), 21(4), 948–962. 10.1210/me.2006-0226.

47. Southall, J., Park, S., Choi, Y., Jeon, H., Ko, C., & Jo, M. (2025). Granulosa cell expression of Fos is critical for regulating ovulatory gene expressions in the mouse ovary. FASEB journal: official publication of the Federation of American Societies for Experimental Biology, 39(4), e70388. 10.1096/fj.202402867R.

48. Liu, Y. X., & Hsueh, A. J. (1986). Synergism between granulosa and theca-interstitial cells in estrogen biosynthesis by gonadotropin-treated rat ovaries: studies on the two-cell, two-gonadotropin hypothesis using steroid antisera. Biology of reproduction, 35(1), 27–36. 10.1095/biolreprod35.1.27.

49. Zhang, Z., He, C., Gao, Y., Zhang, L., Song, Y., Zhu, T., Zhu, K., Lv, D., Wang, J., Tian, X., et al. (2021). α-ketoglutarate delays age-related fertility decline in mammals. Aging cell, 20(2), e13291. 10.1111/acel.13291.

50. Seth, S., Mallik, S., Bhadra, T., & Zhao, Z. (2022). Dimensionality Reduction and Louvain Agglomerative Hierarchical Clustering for Cluster-Specified Frequent Biomarker Discovery in Single-Cell Sequencing Data. Frontiers in genetics, 13, 828479. 10.3389/fgene.2022.828479.

51. Morris, M. E., Meinsohn, M. C., Chauvin, M., Saatcioglu, H. D., Kashiwagi, A., Sicher, N. A., Nguyen, N., Yuan, S., Stavely, R., Hyun, M., et al. (2022). A single-cell atlas of the cycling murine ovary. eLife, 11, e77239. 10.7554/eLife.77239.

52. Pedersen, T., & Peters, H. (1968). Proposal for a classification of oocytes and follicles in the mouse ovary. Journal of reproduction and fertility, 17(3), 555–557. 10.1530/jrf.0.0170555.

