## Supplemental Figures and Table for "FOSL2 regulates gonadotropin-dependent folliculogenesis through feedback amplification of FSH/FSHR signaling"

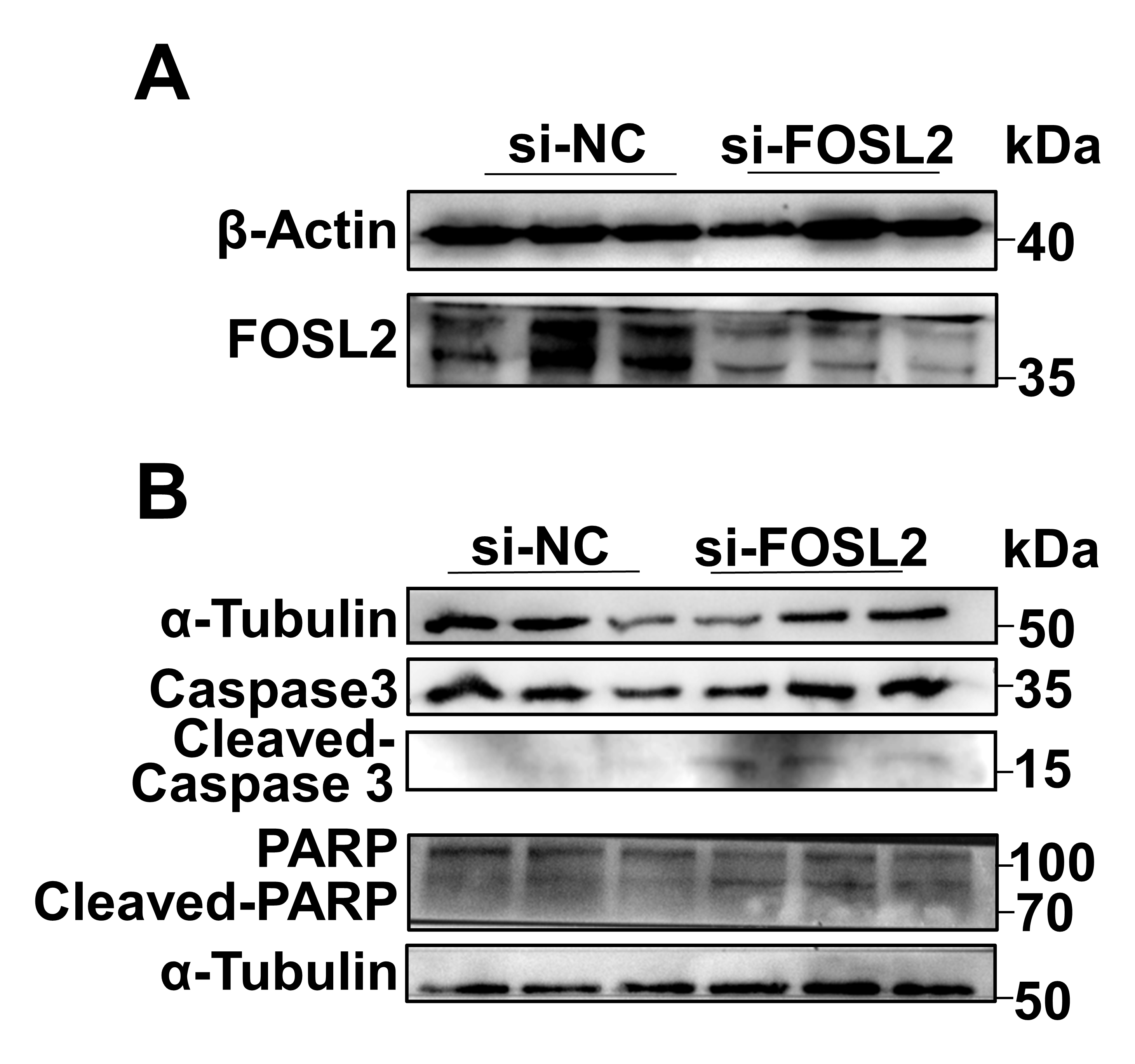

**Figure S1. Efficiency analysis of RNA interference and Knockdown of FOSL2 induces apoptosis** **(related to Figure 2).** (A) Evaluation of FOSL2 knockdown efficiency by Western blotting; n = 3 independent GC samples. Original blots were provided in Figure S6. (B) Changes in pro-apoptotic protein contents following FOSL2 knockdown; n = 3 independent GC samples. Original blots were provided in Figure S6.

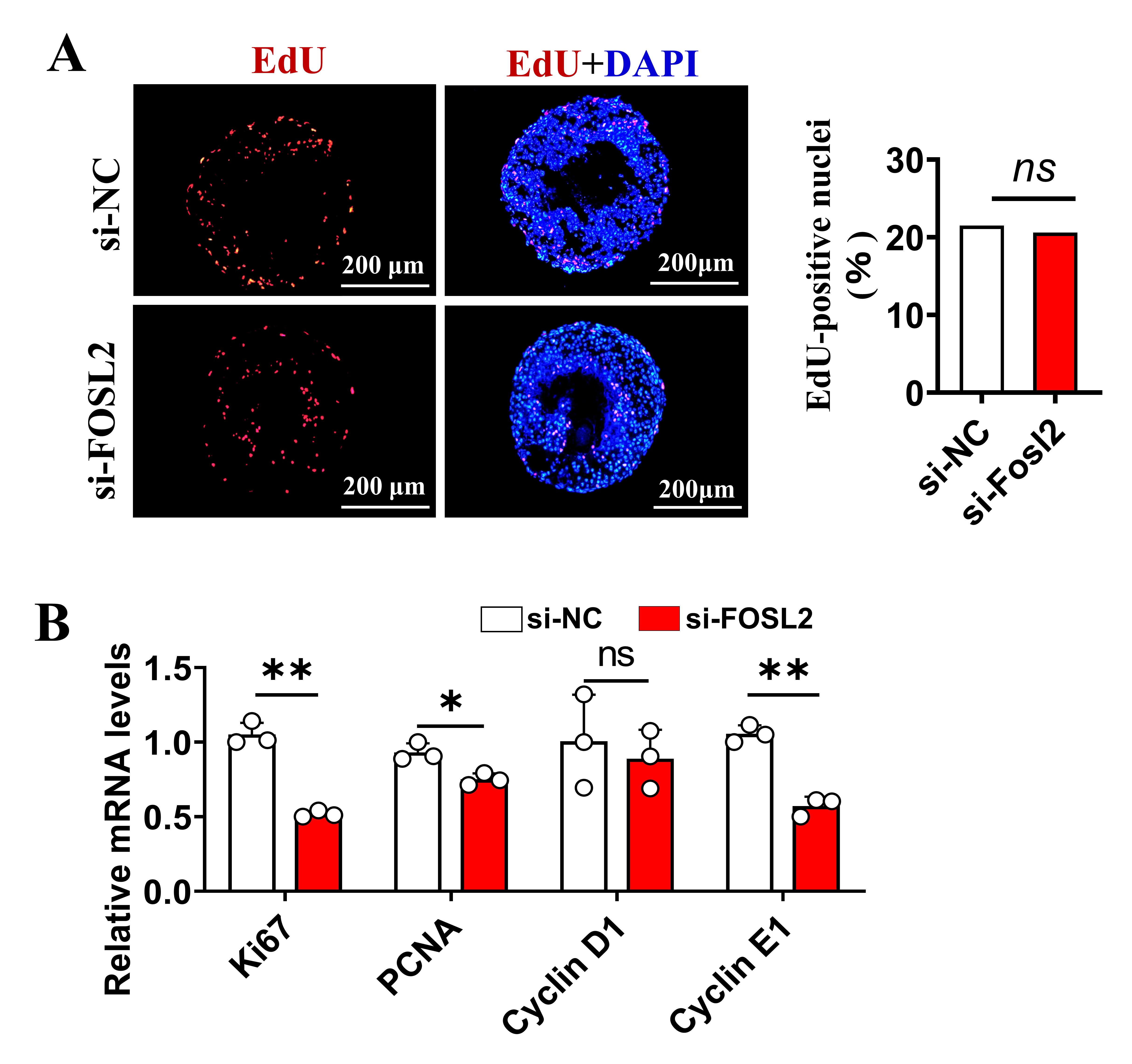

**Figure S2. Knockdown of FOSL2 plays a different role at distinct developmental stages** **(related to Figure 3).** (A) Follicular cell proliferation analysis using the EdU incorporation assay. Left: representative images of EdU staining; right: quantification of EdU-positive nuclei; n = 4 follicles. (B) Expression analysis of proliferation-related genes using qRT-PCR, n = 3 follicles.

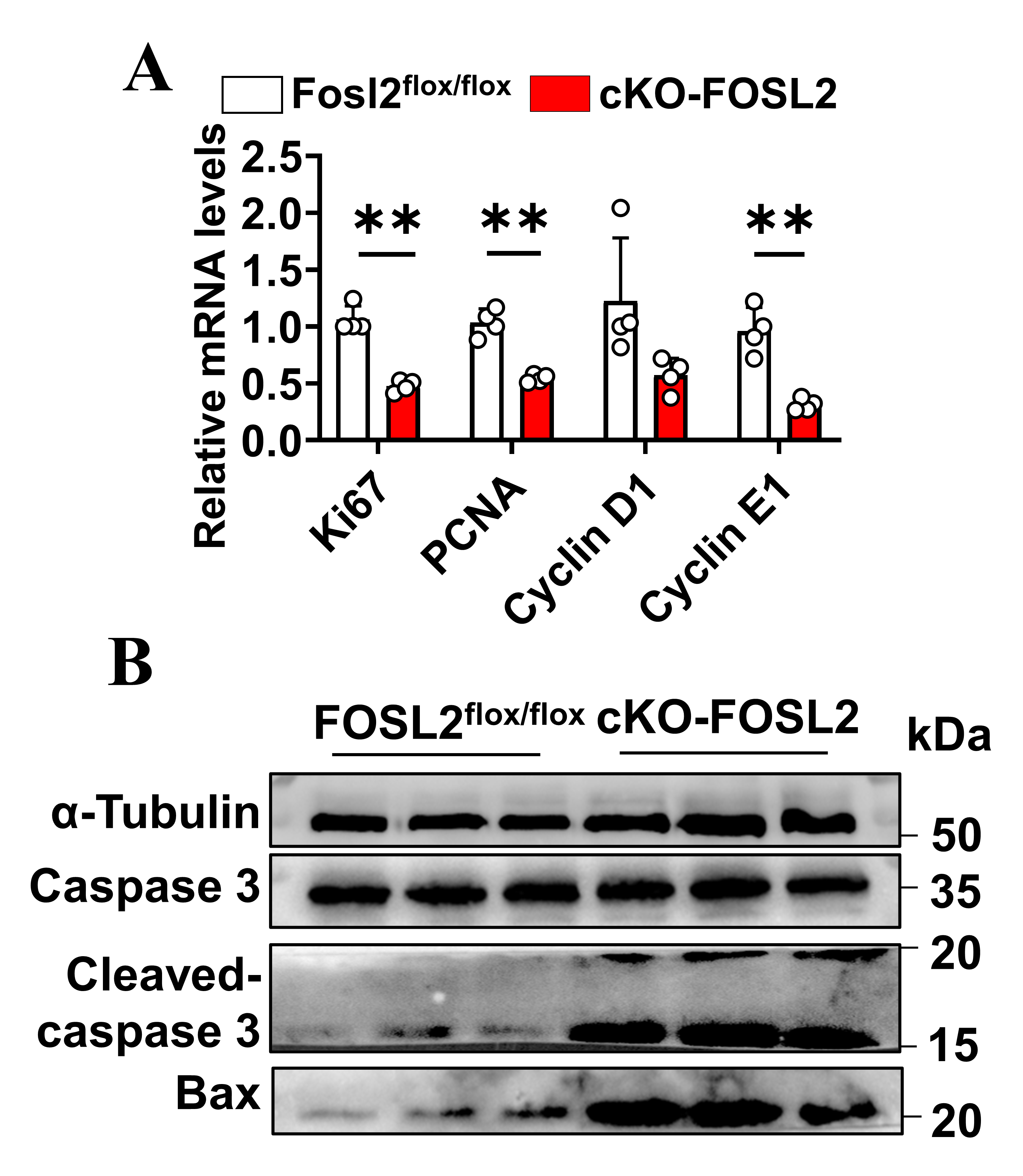

**Figure S3.** **GC-specific FOSL2 knockout** **impaired GC proliferation and induces apoptosis (related to Figure 4).** (A) Expression analysis of proliferation-associated genes using qRT-PCR, n = 4 independent GC samples. (B) Changes in pro-apoptotic protein contents following FOSL2 knockout; n = 3 independent GC samples. GCs were isolated from ovaries 48 hours post-PMSG injection. Original blots were provided in Figure S6.

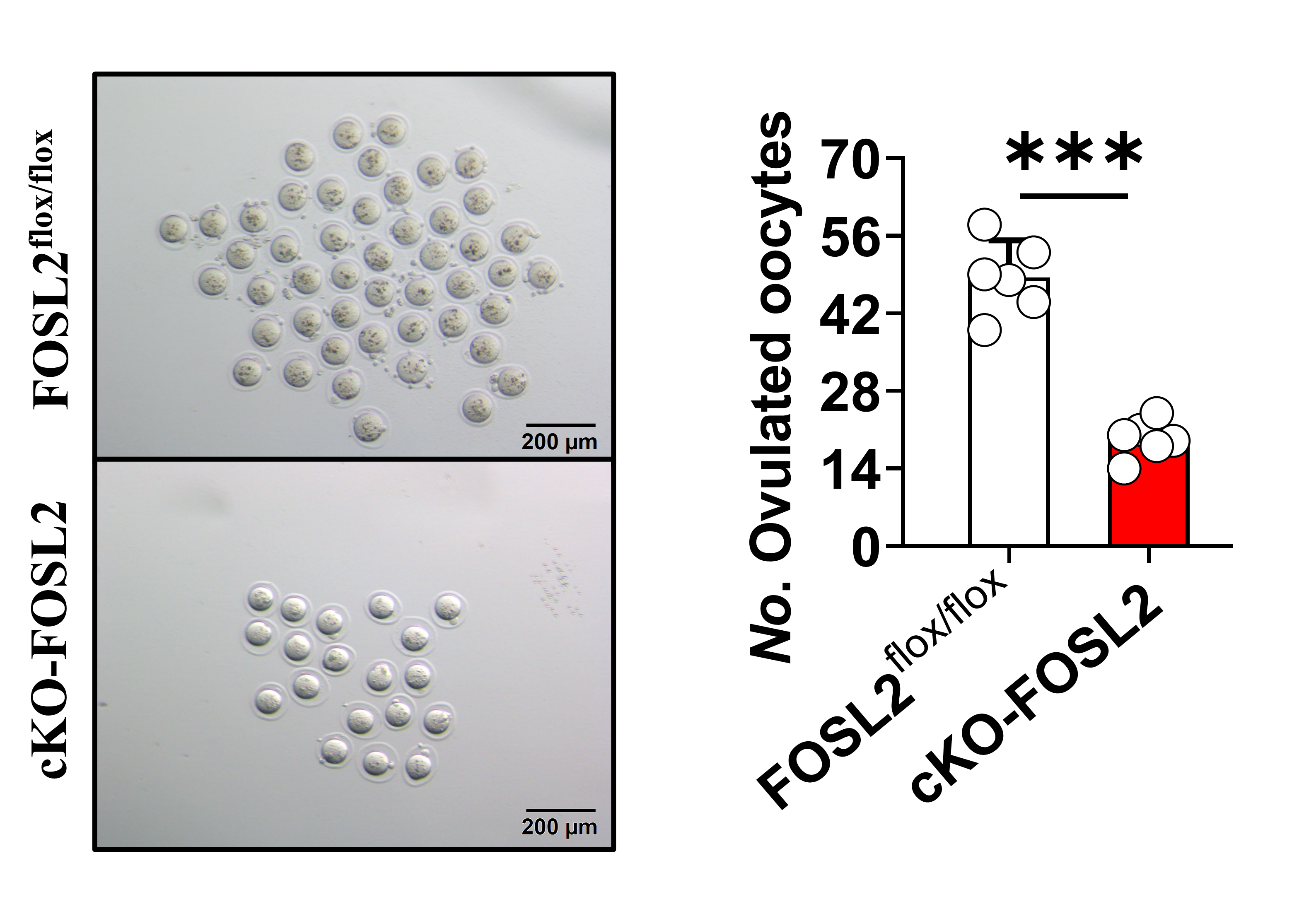

**Figure S4. GC-specific FOSL2 knockout caused significant decrease in the number of ovulated oocytes** **(related to Figure 4).** Analysis of ovulated oocytes. Left: representative images of oocytes released; right: quantification of ovulated oocytes. n = 6 mice.

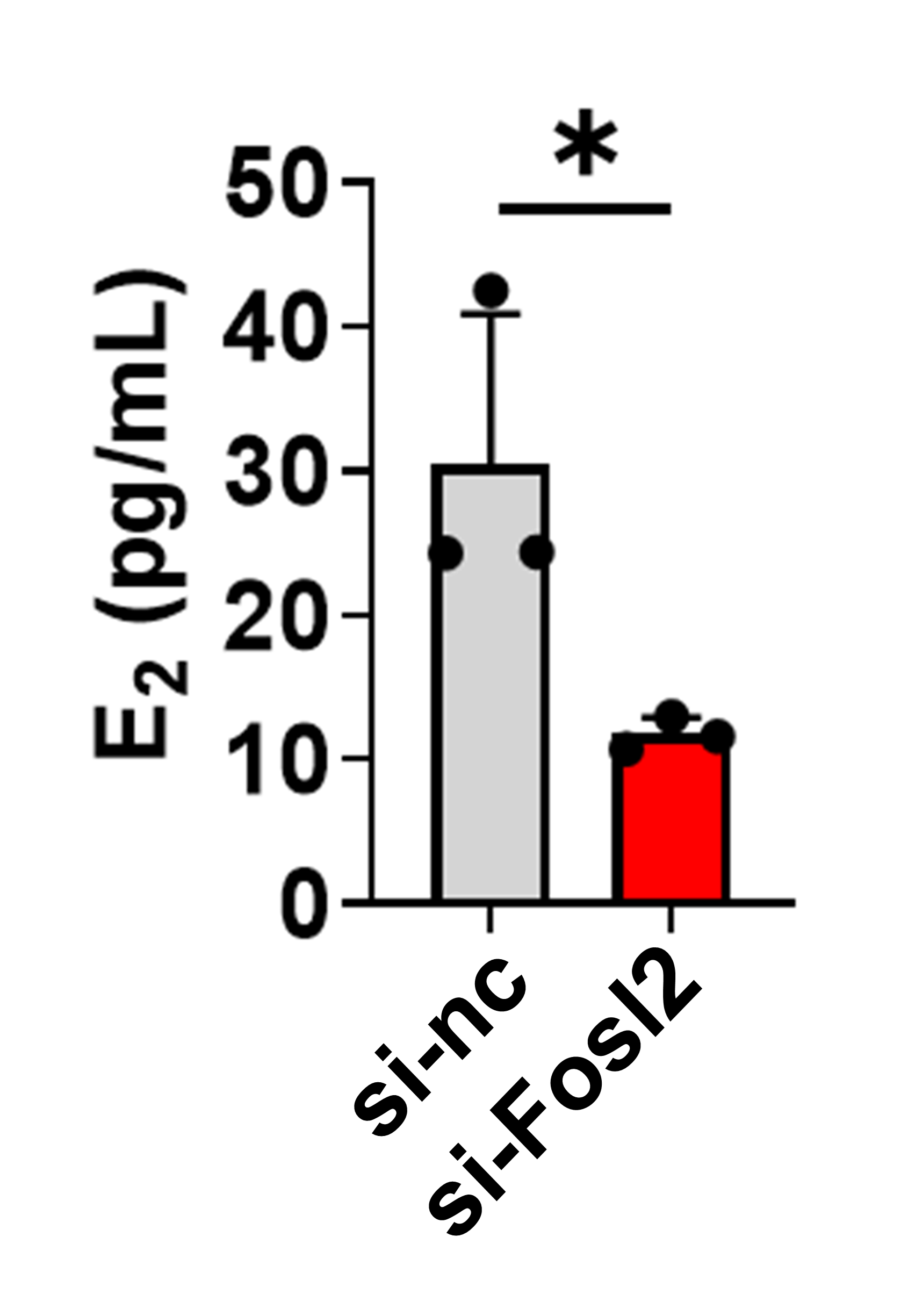

**Figure S5. Knockdown of FOSL2 impaired estrogen biosynthesis of GTH-dependent follicles (related to Figure 5).** Measurement of serum estradiol levels in GTH-dependent follicles following FOSL2 knockdown, n = 3 independent follicular samples.

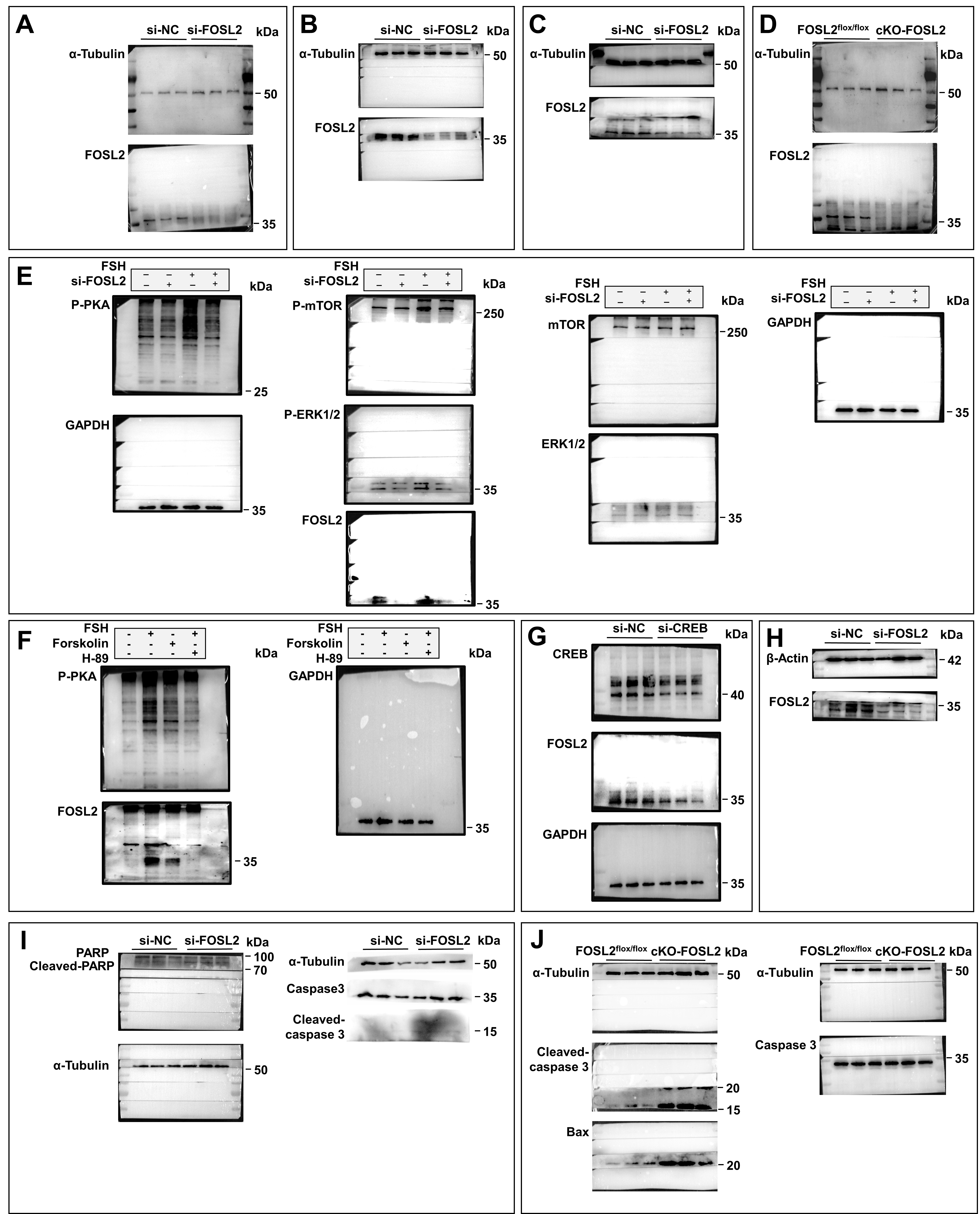
**Figure S6. Full western blots.** (A) Western blot analysis of FOSL2 protein levels in mouse GTH-independent follicles following FOSL2 knockdown (related to Figure 3B). (B) Western blot analysis of FOSL2 protein levels in mouse GTH-dependent follicles following FOSL2 knockdown (related to Figure 3E). (C) Western blot analysis of FOSL2 protein levels in sheep GTH-dependent follicles following FOSL2 knockdown (related to Figure 3K). (D) Western blot analysis of FOSL2 protein levels in cKO mice (related to Figure 4C). (E) FSH-downstream signaling cascades in si-FOSL2 GCs following FSH supplement (related to Figure 5F). (F) Western blot analysis of FOSL2 protein levels in GCs following activation or inhibition of the cAMP-PKA cascade (related to Figure 6C). (G) Western blot analysis of FOSL2 and CREB protein levels in GCs following CREB knockdown (related to Figure 6E). (H) Western blot analysis of FOSL2 protein levels in mouse GCs following FOSL2 knockdown (related to Figure S1A). (I) Western blot analysis of pro-apoptotic protein levels in mouse GCs following FOSL2 knockdown (related to Figure S1B). (J) Western blot analysis of pro-apoptotic protein levels in cKO mice (related to Figure S1B).

**Table S5. The primers used for qPCR and Luciferase reporter**

| Gene | Primer sequence (5'-3') |
| --- | --- |
| *β-Actin-mouse* | Forward: CCAGCCTTCCTTCTTGGGTAT |
|  | Reverse: AGGTCTTTACGGATGTCAACG |
| *Fosl2-mouse* | Forward: GGTAGATATGCCTGGCTCGG |
|  | Reverse: TCATCTCTCCTTCTGCGGCC |
| *Ki67-mouse* | Forward: ATCATTGACCGCTCCTTTAGGT |
|  | Reverse: GCTCGCCTTGATGGTTCCT |
| *Cyclin D1-mouse* | Forward: GCTGGTAGTATGAGGTGCTTGG |
|  | Reverse: CTTTGCAGGACAGATCCCG |
| *Cyclin E1-mouse* | Forward: A GACAAAGCCCAAGCAAAGAA |
|  | Reverse: TGGAGGCAATGGCAGGTT |
| *PCNA-mouse* | Forward: ACCTGCAGAGCATGGACTCG |
|  | Reverse: GCAGCGGTATGTGTCGAAGC |
| *FSHR -mouse* | Forward: TGCTCTAACAGGGTCTTCCTC |
|  | Reverse: TCTCAGTTCAATGGCGTTCCG |
| *CYP11A1-mouse* | Forward: AGGTCCTTCAATGAGATCCCTT |
|  | Reverse: TCCCTGTAAATGGGGCCATAC |
| *β-Actin-sheep* | Forward: CCTGCGGCATTCACGAAACTAC |
|  | Reverse: ACAGCACCCTGTTGGCGTAGAG |
| *Fosl2-sheep* | Forward: CGGGAACTTTGACACCTCGT |
|  | Reverse: TGATGGCGTTGATGGTAGGG |
| *Cyclin B1-sheep* | Forward: TGGCTACTITCCACTTGAGGAT |
|  | Reverse: GGTGACTTGGGCTTACACACA |
| *Cyclin D1-sheep* | Forward: GCTGGTCCTGGTGAACAAAC |
|  | Reverse: CACAGAGGGCAACGAAGGTC |
| *Cyclin E1-sheep* | Forward: AGATGCGCACAACATCCAGA |
|  | Reverse: CAAAGTGAAGAGGCTGCCCA |
| *PCNA-sheep* | Forward: TCTCATGTCTCCTTGGTGCA |
|  | Reverse: GCCAAGGTGTCCGCATTATC |
| *FSHR -sheep* | Forward: ATGCGGTCGAACTGAGGTTT |
|  | Reverse: GCAGGTTGTTGGCCTTTTCA |
| *CYP11A1-sheep* | Forward: GTTTCGCTTTGCCTTTGAGTC |
|  | Reverse: ACAGTTCTGGAGGGAGGTTGA |
| *FOSL2-CYP11a1* | Forward: AGAGGAGGGATGACTCTTGT |
|  | Reverse: ACAAGAGTCATCCCTCCTCT |
| *Mut-FOSL2-CYP11a1* | Forward: AGAGGAGGGAGCACGCTTGT |
|  | Reverse: ACAAGCGTGCTCCCTCCTCT |
| *FOSL2-FSHR* | Forward: CCTTTAGTGGGTCACGTGAC |
|  | Reverse: GTCACGTGACCCACTAAAGG |
| *Mut-FOSL2-FSHR* | Forward: CCTTTAGGCGGCCACGTGAC |
|  | Reverse: GTCACGTGGCCGCCTAAAGG |
| *FOSL2-STAR* | Forward: ACTTCCTGAGTCAATGGCAG |
|  | Reverse: CTGCCATTGACTCAGGAAGT |
| *Mut-FOSL2-Star* | Forward: ACTTCCTGCGTGCATGGCAG |
|  | Reverse: CTGCCATGCACGCAGGAAGT |
| *FOSL2-LHCGR* | TGGCAATGAGCCACTGCACA |
|  | Reverse: TGTGCAGTGGCTCATTGCCA |
| *Mut-FOSL2-LHCGR* | Forward: TGGCAATGCGCTGCTGCACA |
|  | Reverse: TGTGCAGCAGCGCATTGCCA |
| *FOSL2-Inha* | Forward: GCATGTGTGAGTCAGGTCGC |
|  | Reverse: GCGACCTGACTCACACATGC |
| *Mut- FOSL2-Inha* | Forward: GCATGTGCAAGGCAGGTCGC |
|  | Reverse: GCGACCTGCCTTGCACATGC |
| *si-Fosl2-mouse* | Target sequence: gcacttcaaaccttgtcttca |
| *si-Fosl2-sheep* | Target sequence: gtgccgtggtggtgaaaca |
| *si-CREB* | Target sequence: cagcagctcatgcaacatcat |
